# Genetic characterization of stem cell-like cancer cell in the peri-tumoral regions and proliferative lymphocyte in the peripheral blood of patients with glioblastoma

**DOI:** 10.1101/2020.01.06.895987

**Authors:** Runwei Yang, Guanglong Huang, Jinglin Guo, Yaomin Li, Haimin Song, Kaishu Li, Guozhong Yi, Zhiying Lin, Xiran Wang, Sidi Xie, Zhifeng Liu, Xiaofeng Shi, Ke Li, Songtao Qi, Yawei Liu

## Abstract

The glioblastoma (GBM) recurrence rate is high, despite multimodal treatment including surgery, radiotherapy, chemotherapy, and immunotherapy. Most recurrences occur at the resection margin that is located outside the GBM contrast-enhancing region (C region) of magnetic resonance imaging. However, the nature of the GBM cells that lie outside the C region is unclear. We used single-cell RNA sequencing (scRNA-seq) to compare 12 samples taken from inside and outside the C region from four patients with GBM and identified a cluster of GBM cells outside the C region that exhibited fragmented CNVs in chromosomes such as 1, 10, 12 and 19 (herein termed CNV 4). A transcription factor–transcription cofactor interaction network was constructed to uncover the transcriptional regulatory mechanism of these CNV 4 GBM cells that had prognostic significance (p < 0.05). Furthermore, a sub-cluster of these CNV 4 GBM cells possessed stem cell-like properties and had tumorigenic potential. In parallel, we analyzed peripheral blood mononuclear cells (PBMCs) from four patients with GBM, three patients with epilepsy and three healthy volunteers by scRNA-seq. We identified a novel subtype of proliferative immune cells only in patients with GBM. Overall, our findings indicate that CNV 4 cells might contribute to GBM recurrence and proliferative circulating immune cells are specific to patients with GBM. This study sheds light on a neglected aspect of GBM, and opens new avenues to explore GBM recurrence and immunotherapy.

**Significance:** We show that a subset of GBM cells outside the C region possess the properties of stem cell and tumorigenesis potential, which might be responsible for cancer recurrence. Besides, proliferative lymphocytes are specific to the PBMCs of patients with GBM, which could help to develop novel immunotherapy.

## Introduction

Glioblastoma (GBM) is the most common and lethal brain malignancy in adults (1,2). GBM is characterized by contrast enhancement in T1 gadolinium-enhanced magnetic resonance imaging (MRI) with a surrounding non-enhancing region of abnormal T2/fluid-attenuated inversion recovery (FLAIR) signal (3–5). The former region represents the dense cellular GBM core, with neovascularization and blood-brain barrier (BBB) disruption, and the latter region constitutes edematous tissue with infiltrating GBM cells.

Treatment of GBM primarily consists of surgical resection. The contrast-enhancing region (C region) is always resected as much as possible during surgery (5), whereas the non-enhancing region is often left behind after surgery and is the target of postoperative treatment (5). Extensive investigations of primary GBM cells in the C region have thoroughly dissected GBM from the genetic and epigenetic perspective (6–12). However, the conclusions derived from these cells may not be generalizable to the GBM cells outside the C region, as many studies have revealed great differences between them, such as their responses to irradiation, temozolomide, and lomustine (3,4,7,8,13). The highly heterogeneous and infiltrative nature of this cancer means that multimodal treatment including radiotherapy, chemotherapy, and immunotherapy is often unsuccessful (14–18), and the tumors subsequently recur.

Ninety percent of GBM recurrences occur at the resection margin, even after total resection of the contrast-enhancing region (19); and indeed, numerous studies have confirmed that the extent of resection is positively associated with superior outcomes in patients with GBM. For instance, Li et al. showed that additional resection of the surrounding T2/FLAIR abnormality beyond the C region prolonged the median survival time of patients by 5.2 months (20). Lobectomy of GBM in non-eloquent areas also benefits patients in terms of overall survival (OS) and progression-free survival (PFS) (21). These results suggest that a lower number of GBM cells outside the C region is preferable in terms of improving patient outcomes, and emphasize the prognostic and therapeutic value to understanding the nature of residual GBM cells in more detail.

GBM cells outside the C region were first identified by Silbergeld et al. in 1997 (22); since then, an increasing number of investigators have focused on this field and aimed to elucidate the features specific to these cells. However, these studies have yielded conflicting findings regarding the properties of GBM cells outside and inside the C region. For instance, Silbergeld et al. found no difference in motility between GBM cells isolated from the C region and outside the C region (22), while others demonstrated more invasive features of GBM cells outside the C region than inside the C region (13,23). Similarly, some researchers have shown that GBM cells outside the C region proliferate faster than those derived from the C region (13,22), whereas others have reported the opposite (24,25). These seemingly contradictory results might be partially explained by the heterogeneity of GBM cells, which has been extensively revealed by multiomics studies (6,8–11,26). Furthermore, these results indicate that our current understanding of GBM cells outside the C region is incomplete, which might impede the efficacy of postoperative treatment.

In this study, we aimed to investigate the genetic characterization of GBM cells both inside and outside the C region. To do so, we performed single-cell RNA sequencing (scRNA-seq) on samples from the C region, peri-enhancement edematous region (P region), and peri-edema normal region (N region) from four patients with GBM. We found a unique subset of GBM cells with the properties of cancer stem cell in the P and N regions that might be responsible for recurrence after surgery. To explore whether GBM cells exist in the circulation and to characterize the systemic immune response to GBM, we sampled peripheral blood mononuclear cells (PBMCs) from the four patients with GBM, three patients with epilepsy and three healthy volunteers and analyzed the cells by scRNA-seq. Remarkably, proliferative immune cells were discovered in the PBMCs of patients with GBM but not in patients with epilepsy or in healthy volunteers. Our findings suggest that a specific subset of GBM cells within the resection margin could be targeted for treatment to prevent tumor recurrence.

## RESULTS

### GBM cells show high heterogeneity with respect to copy number variations (CNVs) and a subset of them possess atypical pattern

To investigate the characteristics of primary GBM cells located inside and outside the C region, we collected samples of the C, P, and N regions from four patients with GBM, under MRI guidance (Figure 1A, Supplementary Table S1). Hematoxylin and eosin (HE) staining and immunohistochemistry (IHC) showed that the C region contained the histological hallmarks of GBM in terms of the extent of microvasculature, proliferation and tissue necrosis, as well as a high ratio of KI67-positive cells. Conversely, the P and N regions showed no typical GBM features but resembled, to some extent, normal brain tissue (Figure S1A, B). After preparing a single-cell suspension for each sample, we profiled these twelve GBM samples and four corresponding PBMC samples from four patients with GBM mentioned above on a 10× Chromium 3’ Single Cell Platform to determine their transcriptome status. In total, 72,620 cells passed our quality control filters, including 48,770 cells from GBM samples and 23,850 cells from PBMC samples (Figure 1B, C).

**Figure 1.**
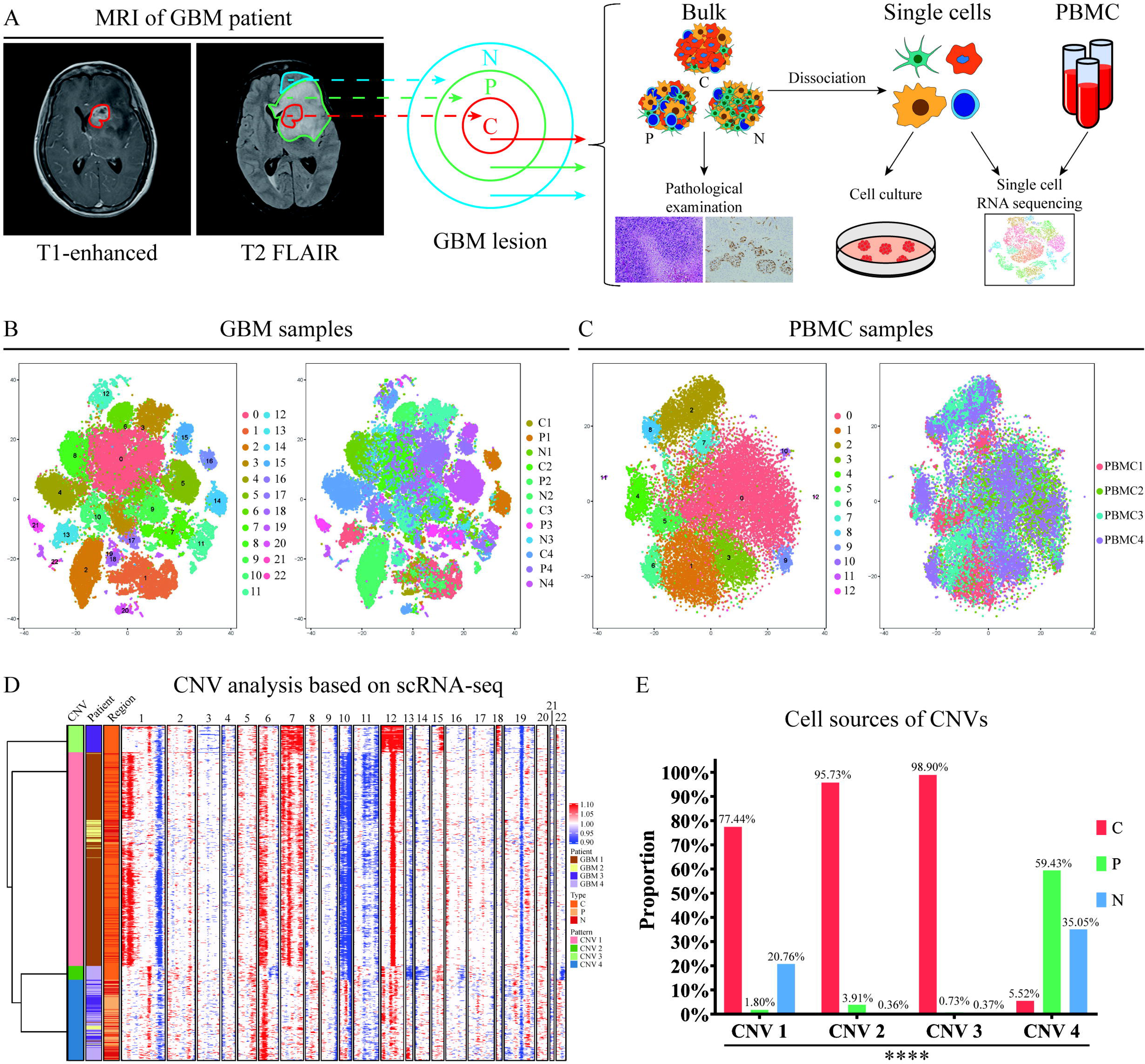
**A**, The study workflow. Samples from the central contrast-enhancing region (C region), peri-enhancement edematous region (P region), and peri-edema normal region (N region) were collected separately under the guidance of MRI. Then, these samples were used for the pathological examination, scRNA-seq analysis, and primary culture of GBM cells. PBMCs of the corresponding patients were also collected for scRNA-seq analysis. **B**, t-SNE plot of 48,770 cells from twelve tissue samples of four patients with GBM. The clustering results of these cells (left) and the sample origins of the different clusters (right) are shown. **C**, A t-SNE plot of the 23,850 cells from four PBMC samples of the corresponding patients with GBM. The clustering results of these cells (left) and the sample origins of the different clusters (right) are shown. **D**, Classification of CNV patterns inferred from the transcriptomic profiles of 6,715 GBM cells. CNV patterns 1, 2, and 3 were mainly patient-specific, while CNV pattern 4 contained GBM cells from four patients. **E**, Spatial distribution of GBM cells with different CNV patterns in the C, P and N regions. Chi-square test, P < 0.0001.

We first classified the cells as being either malignant or nonmalignant cells as follows. First, we divided the cells into 23 clusters according to the Seurat “FindClusters” function, each of which represented a transcriptome-distinct cell group (Figure 1B, C). Then, we identified the differentially expressed genes (DEGs) for each cluster by comparing to the remaining clusters. We used these cell type-specific DEGs to infer the cell identities, and then performed a Gene Ontology (GO) enrichment analysis as confirmation. We identified oligodendrocytes, macrophages, T cells and putative GBM cells in the GBM samples (Figure 1B), and T cells, B cells, and monocytes in the PBMC samples (Figure 1C).

For the 6,715 putative GBM cells that could not be classified as any known cell type, we inferred the CNVs from transcriptomic information using the moving average method(6,10,27,28). Using this algorithm, we identified large-scale amplifications and deletions in putative GBM cells, such as chromosome 7 gain and chromosome 10 loss, which are genetic hallmarks of most GBM cells (Figure 1D). We observed a high degree of heterogeneity with respect to the CNVs among the GBM cells, which is consistent with previous reports (6,10,27). Based on the CNV profile, we could classify the GBM cells into four CNV patterns (Figure 1D). GBM cells of CNV pattern 1 had typical CNV hallmarks of chromosome 7 amplification and chromosome 10 deletion, as well as variations in chromosomes 1, 6, 11, 12 and 19. GBM cells of CNV pattern 2 featured the regional loss of chromosomes 6, 10, 13, 14, 16, 19 and 22, while those of CNV pattern 3 showed amplifications of chromosomes 7, 12, 15 and 18. GBM cells of CNV pattern 4 had fragmentary variations in chromosomes 1, 6, 10, 11, 12, 13 and 19, but these were not as typical as the former three patterns. Notably, most GBM cells of CNV patterns 1, 2 and 3 (77.44%, 95.73%, and 98.90%, respectively), whose CNVs were similar to classical GBM hallmarks, were derived from the C region, whereas GBM cells of CNV pattern 4 were found mainly in the P and N regions (59.43% and 35.05%, respectively) (chi-square test, P < 0.0001) (Figure 1E). These data suggest that GBM cells in C region and outside of C region have different CNV patterns, which might be employed by GBM cells for malignant transformation and infiltrative progression.

### CNV patterns contribute to the transcriptomic status of GBM cells

Next, we divided GBM cells into 11 transcriptomic sub-clusters by the Seurat “FindClusters” function (Figure 2A). Interestingly, sub-cluster 3 accounted for 93.93% of CNV pattern 3, and sub-cluster 5 accounted for 94.31% of CNV pattern 2, while CNV patterns 1 and 4 were comprised of diverse sub-clusters in relatively balanced proportions (Figure 2B). From another perspective, we found that most transcriptomic sub-clusters were derived from multiple CNV patterns (Figure S2A, B). In other words, there was no strict correspondence between CNV patterns and transcriptomic sub-clusters, and GBM cells with different CNV patterns tended to resemble each other at the transcriptome level to a certain extent. To understand the relationship among GBM cells with different CNV patterns, we analyzed the developmental trajectory of these GBM cells. By Monocle2 algorithm, which is designed for inferring the developmental relationship of cells (29,30), GBM cells were divided into three states, representing three major transcriptomic statuses (Figure 2C). We found that the GBM cells of CNV patterns 1 and 4 were located across three states, while GBM cells of CNV patterns 2 and 3 were mainly isolated in state 2 (Figure 2C). Interestingly, the pseudotime analysis results of Monocle2 algorithm suggested potential transitions from CNV pattern 4 to CNV patterns 1, 2, and 3 or from CNV pattern 1 to CNV patterns 2 and 3 (Figure 2C, Figure S2C).

**Figure 2.**
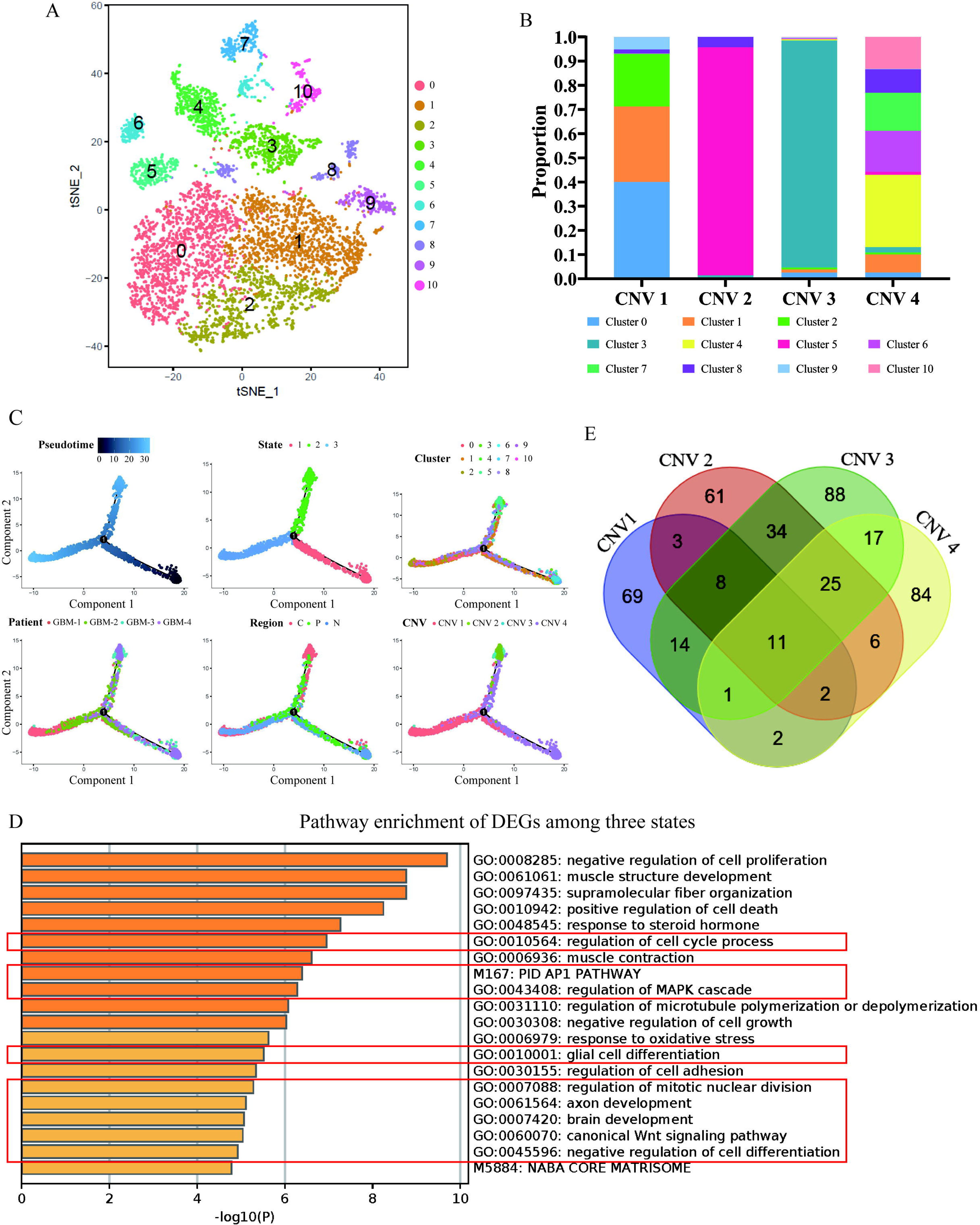
**A**, A t-SNE plot of 6,715 GBM cells. **B**, The cumulative percentage of the different transcriptomic cluster compositions of the four CNV patterns. **C**, The developmental trajectory of all GBM cells inferred by the Monocle2 algorithm. **D**, Pathway enrichment results of the DEGs among the three states. The pathways involved in the regulation of gene expression are marked by red rectangles. **E**, Venn diagram showing the intersection of the DEGs of the four CNV patterns.

Pathway enrichment analysis of the DEGs among different states revealed disturbances in many pathways involved in gene expression regulation, such as the AP1 pathway, glial cell differentiation, the regulation of mitotic nuclear division, and the canonical Wnt signaling pathway (Figure 2D). Notably, 23% of the top 100 DEGs, including *SOX2*, *SOX4*, *SOX11*, *ETV1*, and *CBX3*, were transcription factors (TFs) and transcription cofactors (TCFs) with known roles in GBM (31–37) (Figure S2D and Supplementary Table S2). In addition, an intersection analysis of the DEGs of the four CNV patterns showed that only 11 genes were commonly upregulated. Most of the other DEGs were CNV pattern-specific (Figure 2E). Consistent with these findings, the pathway enrichment analysis of CNV pattern 1-specific DEGs and CNV pattern 4-specific DEGs indicated differences in the transcriptional regulation of gene expression (Figure S2E, F). Taken together, these findings suggest that CNV patterns and transcriptional regulation cooperate to shape GBM transcriptomic states.

### Transcriptional regulatory networks orchestrate the GBM cell transcriptome and correlate with patient outcomes

To investigate how GBM cells are transcriptionally regulated, we focused on the TFs and TCFs identified as described above. We first established a strategy to construct the TF–TCF interaction network for the four patterns (Figure 3A). Briefly, we selected TFs and TCFs in the DEGs of the four CNV patterns. TFs and TCFs cooperate to exert their functions; therefore, we examined their interactions according to the Search Tool for the Retrieval of Interacting Genes/Proteins (STRING) database. To obtain confident interactions, we only considered those derived from curated databases and experiments and with combined scores >0.9 (Figures S3, S4, S5, and S6). After sorting by the interaction degree of each TF, we chose the top three TFs as hub TFs to further analyze the differentially expressed target genes and to construct the TF–TCF-target regulatory networks (Figure 3B, Figures S7, S8, S9, and S10). We found that JUN was implicated in all four CNV patterns, MYC was involved in CNV patterns 2 and 4, TP53 and E2F1 were specific to CNV pattern 1, SMAD2 was specific to CNV pattern 3, and FOS was specific to CNV pattern 4 (Figure 3B, C). Additionally, we identified differentially expressed TFs and TCFs with fold changes (FCs) >2 compared to normal cells.

**Figure 3.**
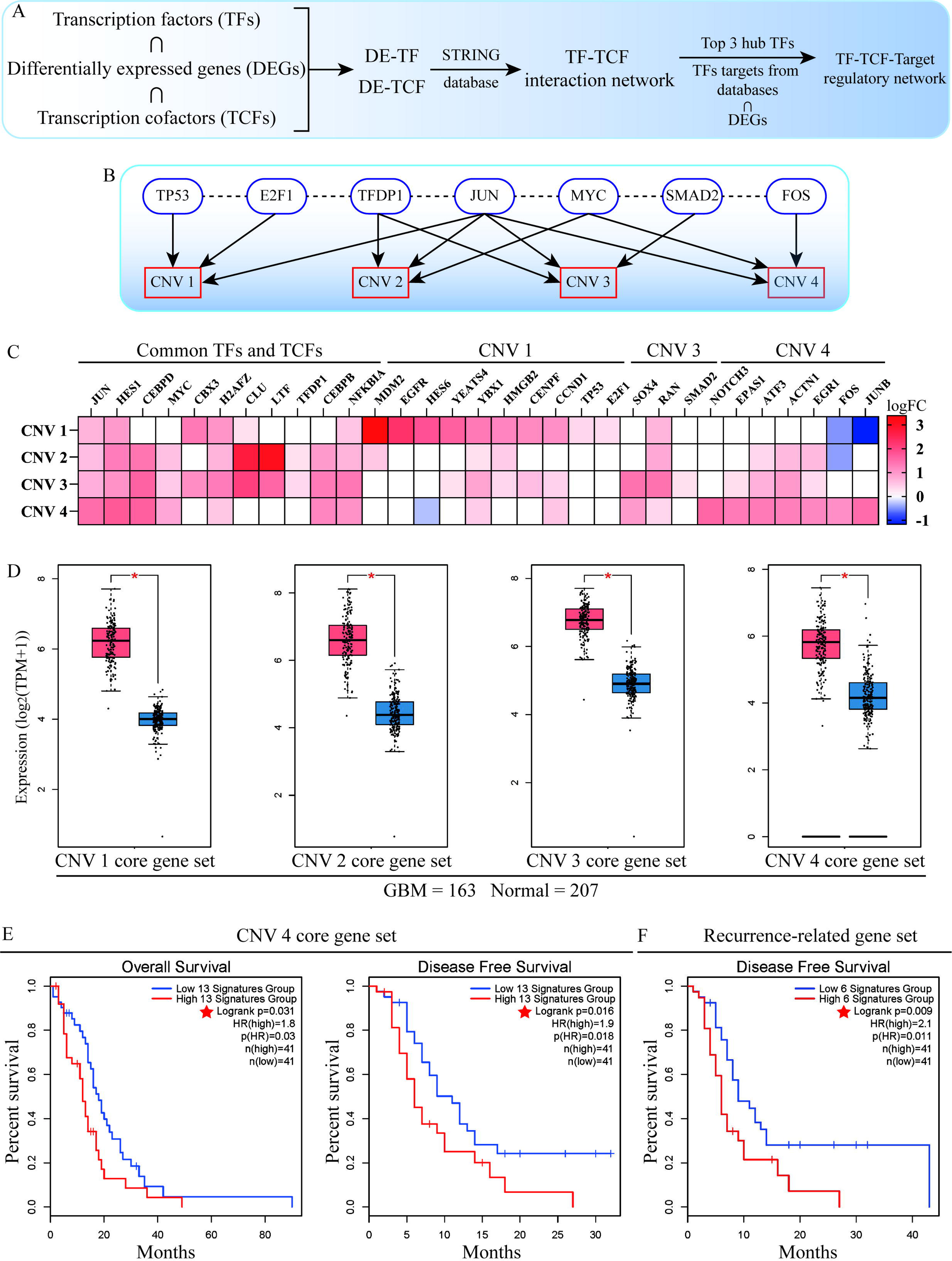
**A**, The strategy used to construct the TF–TCF-target regulatory network. **B**, The top three TFs of the four CNV patterns with the highest interaction degrees (interacting neighbors). **C**, Heatmap of the core gene sets of the TF-TCF interaction networks of the four CNV patterns. The color gradation represents the log(FC) of the DEGs. The common TFs and TCFs in different CNV patterns and CNV pattern-specific TFs/TCFs are arranged from left to right. **D**, Expression analysis of the four CNV pattern core gene sets based on The Cancer Genome Atlas (TCGA) and the Genotype-Tissue Expression (GTEx) databases. **E**, Prognostic value of the CNV pattern 4 core gene set in patients with GBM included in the TCGA database. The expression value of the gene set was defined as the average value of all the genes in the corresponding gene set. Patients with expression values exceeding the 75th percentile were grouped into the high signature group, and those with expression values less than the 25th percentile were grouped into the low signature group. **F**, The prognostic value of the recurrence-related gene set in patients with GBM from the TCGA database.

By combining TFs and TCFs with high interaction degrees and variations, we defined the core TF and TCF gene sets for the four CNV patterns (Figure 3C, Figure S11A). The subnetworks of the core gene sets comprised 61.54%, 41.41%, 56.95%, and 58.56% of the TF–TCF interaction networks for CNV patterns 1, 2, 3 and 4, respectively (Figures S3, S4, S5, and S6). These CNV-specific core gene sets were, however, derived from the scRNA-seq analysis of only four patients with GBM. Therefore, to determine whether they were applicable in a larger cohort, we analyzed data from The Cancer Genome Atlas (TCGA) and Genotype-Tissue Expression (GTEx) databases and found that these gene sets were also more highly expressed in GBM tissue than in normal brain tissue (Figure 3D, Figure S11B, C, D, E, F, G, H). Interestingly, the CNV pattern 4–specific TFs *JUNB* and *FOS* showed the opposite expression patterns to CNV pattern 1 (Figure 3C, Figure S12A, B, C, D). Besides, we found that these TFs have different roles depending on the cancer context. For example, both *JUNB* and *FOS* are upregulated in GBM, acute myelocytic leukemia (LAML), and pancreatic adenocarcinoma (PAAD), but are downregulated in some other malignancies, such as bladder urothelial carcinoma (BLCA), lung adenocarcinoma (LUAD), and ovarian serous cystadenocarcinoma (OV) (Figure S12A, B, C, and D). These results indicated that TFs might play different roles in different GBM cells.

We next used the CNV gene sets to analyze survival in the TCGA GBM cohort. Interestingly, only the core gene set of CNV pattern 4 was negatively correlated with both OS and disease-free survival (DFS), while the core gene sets of other CNV patterns showed little discernible correlation (Figure 3E, Figure S13A, B). We thus compared the expression of the CNV pattern 4 core gene set between primary GBM and recurrent GBM to gain a genetic perspective on the potential mechanisms underlying GBM recurrence. We found that *JUNB*, *ATF3*, *ACTN1*, *CEBPB*, *EGR1*, and *NFKBIA* were significantly highly expressed in recurrent GBM and positively associated with each other (Figure S13C, D). As expected, these recurrence-related genes (RRGs) were negatively associated with DFS (Figure 3F). These results suggested that CNV pattern 4 GBM cells might play a vital role in cancer recurrence.

### A subset of CNV pattern 4 GBM cells possess stem-cell properties

Much research has revealed that GBM recurrence after surgery often initiates at the peri-tumoral regions (P and N regions). Earlier analysis showed that the core gene set of CNV 4 GBM cells was closely associated with cancer recurrence in patients with GBM. Therefore, we aimed to determine whether CNV 4 GBM cells in the P and N regions have gliomagenesis ability. An analysis of the CNV compositions of each GBM sample showed that >94% of GBM cells in the P and N regions of two patients with GBM (recorded as GBM-3 and GBM-4 in this study) belonged to CNV pattern 4 (Figure 4A, Figure S14A). Notably, 100% of GBM cells in the N region of GBM-3 were clustered into CNV pattern 4 (Figure 4A). We then established sphere cultures of the primary GBM cells derived from samples of C, P and N regions of GBM-3 to test their potential of tumorigenesis. As expected, GBM cells from the C, P and N regions formed spheres and expressed the stem cell marker CD133 (Figure 4B), indicating that CNV 4 GBM cells have tumorigenic potential (31,38–40).

**Figure 4.**
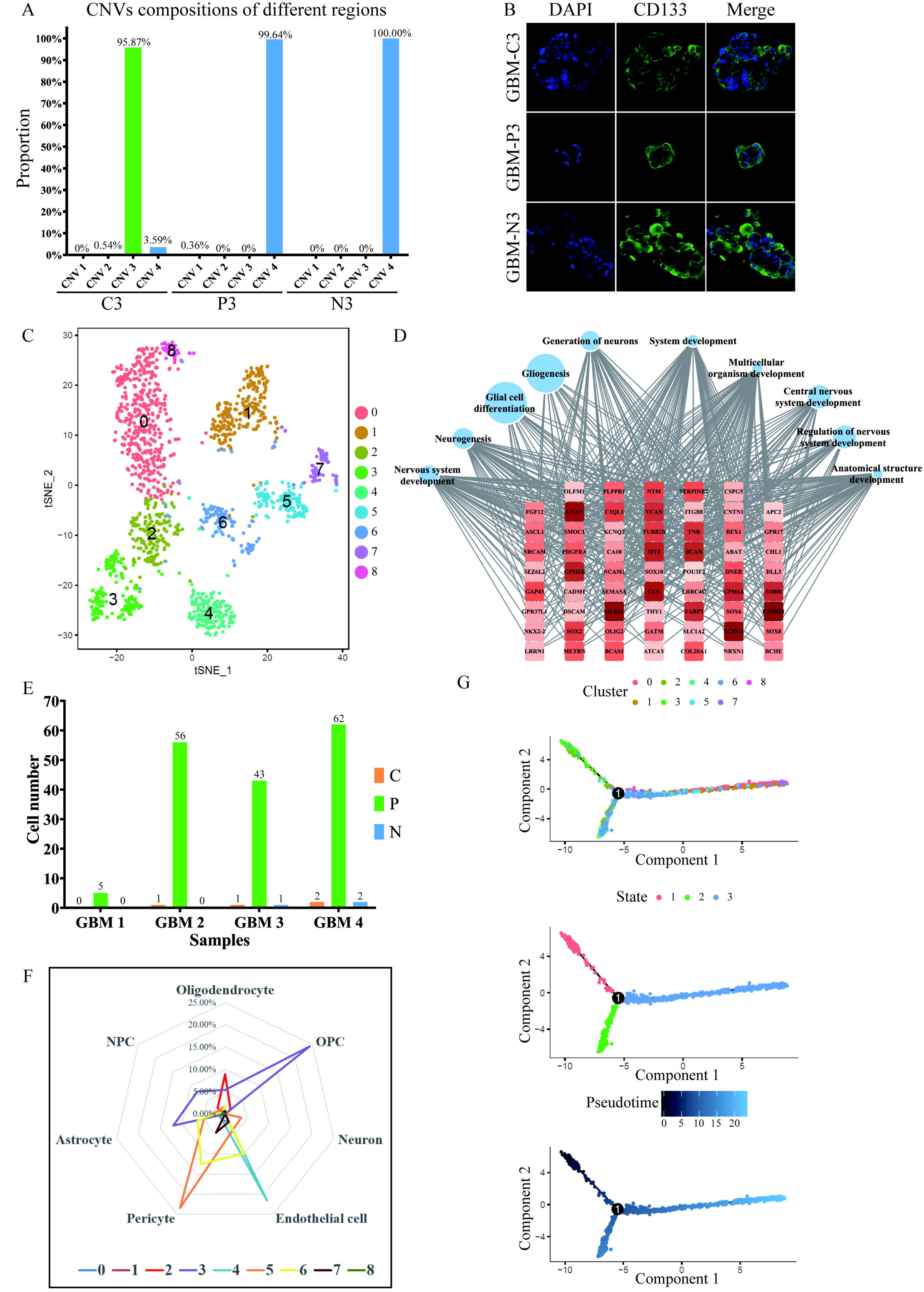
**A**, The CNV pattern compositions of different samples from GBM-3 patients. **B**, Immunofluorescent labeling of GBM spheres for glioma stem cell (GSC) marker CD133 expression. These GBM cells were derived from the C, P and N regions of GBM-3. **C**, t-SNE plot of 1,612 CNV pattern 4 GBM cells. **D**, Results of the GO enrichment analysis of sub-cluster 3 differentially expressed genes (DEGs). The DEGs are listed in rounded rectangles. The FCs of the DEGs are reflected by the color gradation. A larger FC corresponds to a deeper color. The top 10 enriched biological processes (BPs) with the lowest false discovery rates (FDRs) are listed from left to right in circles. A higher level of fold enrichment corresponds to a larger circle. The black lines depict connections between DEGs and BPs. **E**, The spatial distribution of sub-cluster 3 among 12 GBM samples. **F**, The extent of overlap between the DEGs of different sub-clusters and different cell types. **G**, The developmental trajectory of CNV pattern 4 GBM cells inferred by the Monocle2 algorithm.

To characterize the properties of CNV 4 GBM cells in more detail, we furtherly analyzed their sub-clusters. As shown in figure 4C, altogether 1,612 GBM cells of CNV 4 were divided into nine sub-clusters (Figure 4C). Interestingly, GO enrichment analysis of the sub-cluster DEGs revealed that the cells comprising sub-cluster 3 were involved in neurogenesis, gliogenesis, and stem cell differentiation (Figure 4D, Figure S14B) and were mainly located in the P region (166/173) (Figure 4E). The GBM stem-like cell marker *SOX2* was highly expressed in this sub-cluster (Figure 4D), and IHC confirmed that SOX2-positive and Nestin-positive cells existed in both the P and N regions (Figure S15A). Consistent with our earlier analysis (Supplementary Figure 2F), some of the other sub-cluster DEGs were enriched in the biological processes associated with angiogenesis and muscle structure development.

These results brought to mind the GBM stem cells (GCSs) and vasculogenic mimicry (VM) that could be formed by CD133-positive GCSs (41,42). We then evaluated the resemblance of sub-clusters to endothelial cells, pericytes, and other central nervous system cell types, including astrocytes, oligodendrocytes, oligodendrocyte progenitor cells (OPCs), neurons, and neural progenitor cells (NPCs) (the cell type-specific gene sets are listed in Supplementary Table S3) (43,44). As expected, sub-cluster 3 expressed markers of astrocytes, oligodendrocytes, OPCs and NPCs simultaneously (11.97%, 5.36%, 24.24%, and 8.03%, respectively), while the other sub-cluster DEGs overlapped with endothelial cell or pericyte marker genes to different extents (from 0% to 21.57% and from 0% to 23.44%, respectively) (Figure 4F). We also established stemness, endothelial cell and pericyte scores according to the average expression of the relevant marker genes to estimate the properties of CNV pattern 4 GBM cells. These three scores revealed continuums rather than separate distributions of stemness and differentiations towards endothelial cells or pericytes (Figure S14C). Next, we performed a pseudotime analysis using the Monocle2 algorithm to draw the developmental trajectories of CNV pattern 4 GBM cells based on their transcriptomic similarities (29,30). Consistent with the former result, sub-cluster 3 cells were located at the initial state, and the other sub-clusters were distributed along the later states (Figure 4G, Figure S14D). Collectively, these results revealed that the sub-cluster 3 of CNV 4 GBM cells possessed stemness properties and might be able to differentiate into endothelial cells and/or pericytes.

### Sub-clusters of intracranial immune cells and PBMCs from patients with GBM show abnormal gene-expression pattern

Previous study showed that the changes of immune microenvironment were involved in the gene-expression pattern, treatment resistance, and recurrence of GBM (8). Therefore, we additionally analyzed the intracranial immune cells, which constitute the largest proportion of sequenced cells in GBM tissue samples. CNV analysis demonstrated that these cells had a uniform expression pattern but possessed no known hallmarks of GBM cells (Figure 5A). Sub-clusters 12 and 17 interested us, as they exhibited peculiar gene-expression pattern (Figure 5B, Figure S16A). Both of these sub-clusters expressed the marker genes of microglia/macrophages, namely CSF1R, CD14, CD68, AIF1, and HLA-DRB1 (Figure 5B). Moreover, sub-cluster 12 showed simultaneously high expression of EGFR, PTPRZ1, SOX2, SLC35E3, and MDM2, known GBM-associated genes, while sub-cluster 17 was characterized by the expression of the proliferation markers KIAA0101, BIRC5, MKI67, NUSAP1, and TOP2A (Figure 5B). Interestingly, GO enrichment analysis of the DEGs indicated that sub-cluster 12 was associated with nervous system development (Figure 5C), and sub-cluster 17 was enriched in the biological processes of the cell cycle (Figure 5D). However, when compared to GBM cells, these cells exhibited the characteristics of immune cells (Figure 5E, Figure S16B).

**Figure 5.**
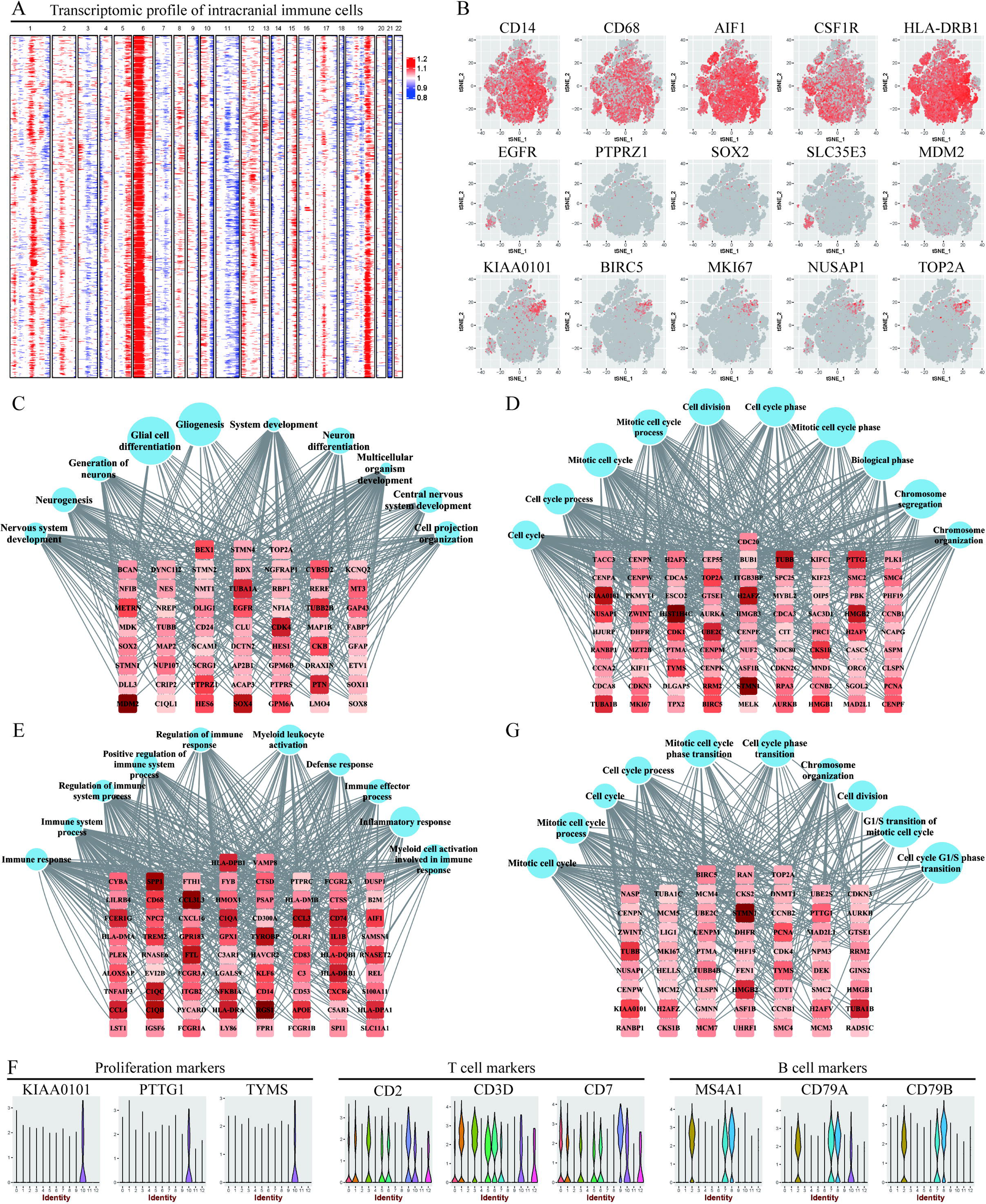
**A**, Genomic profiling data from 23,850 immune cells in 12 GBM tissue samples. **B**, Heatmaps showing markers of microglia/macrophages, malignant cells, and proliferative cells. **C** and **D**, The GO enrichment analysis of subcluster 12 and 17 DEGs compared to the remainder of the other sub-clusters. **E**, The GO enrichment analysis of sub-cluster 12 DEGs compared to GBM cells. **F**, The expression of markers of proliferative cells, T cells, and B cells in the PBMCs of patients with GBM. **G**, The GO enrichment analysis of PBMC sub-cluster 10 DEGs compared to the remainder of the other sub-clusters.

We noticed that leukocyte-associated processes were common to both sub-clusters 12 and 17 (Figure 5E, Figure S16B); therefore, we wanted to determine whether these cells were derived from the peripheral blood. Besides, we wanted to understand whether the CNV pattern 4 GBM cells outside the C region would existed in the circulation of patients with GBM. As such, we analyzed PBMC samples obtained from these four patients with GBM before surgery by scRNA-seq. Interestingly, we detected a cluster of proliferative cells (cluster 10, 125/23850) in their PBMCs (Figure 1F, Figure 5F, G). However, we did not find cells simultaneously expressing both monocyte markers and GBM-associated markers (Figure S16D, E). These proliferative cells were composed of T cells and B cells, but not monocytes (Figure 5F, Figure S16E). When compared to GBM cells, these cells also showed an involvement in immune system-related processes (Figure S16F). Notably, we found that the proliferative T cells simultaneously expressed perforin and granzyme-related genes such as PRF1, GZMA, GZMB, GZMM, and GNLY, while proliferative B cells showed the upregulated expression of immunoglobulin-related genes such as IGKC, IGKV1-12, and JCHAIN, indicative of plasma cells (Figure S16F). These transcriptomic profiles suggested that they were highly responsive to GBM.

To investigate whether these cells also exist in non-tumor patients or healthy ones, we also examined PBMC samples collected from three healthy volunteers and three non-tumor patients (patients with epilepsy) by scRNA-seq. We did not detect any clusters of cells highly expressing proliferation markers in the PBMCs of healthy volunteers or patients with epilepsy (Figure S16G, H), suggesting that these proliferative lymphocytes were specific to the PBMCs from patients with GBM.

## DISCUSSION

In this study, we identified four CNV patterns in twelve GBM samples by scRNA-seq. CNV pattern 4 showed a lower degree of CNVs than the other three patterns. Interestingly, > 90% of CNV pattern 4 GBM cells were distributed in the P and N regions of four patients with GBM, while the other CNV pattern GBM cells were found mainly in the C region of particular patients, consistent with previous studies (10,24). Furthermore, pseudotime analysis revealed a transitional trend from CNV pattern 4 to other CNV patterns. The homogeneous distribution and moderate CNV state of CNV pattern 4 GBM cells suggest the common mechanism of infiltration among highly heterogeneous GBM cells and the possibility of CNV pattern 4 as the prodromal and transitional status of fully transformed malignant cells.

There is likely a complicated course of evolution leading from a normal diploid karyotype to the diverse CNVs detected in the present study, and from normal precursor cells to full-fledged malignant cells. Some researchers have proposed that GBM is derived from the malignant transformation of NPCs/neural stem cells (NSCs) (45–47). GBM precursors are normal diploid cells, whereas GBM cells are characterized by trisomy 7 and monosomy 10, as well as other notable CNVs (amplification of EGFR, PDGFRA, and MET and deletion of CNKN2A, PTEN, etc.) (48). Neftel et al. inferred heterogeneous CNV patterns of GBM cells based on transcriptomes (both intertumoral and intratumoral) (6). Notably, the GBM hallmarks of chromosome 7 amplification and chromosome 10 deletion were not detected in the GBM cells of some adult and most pediatric patients (6,27). Distinct from most typical GBM cells, there are fewer CNVs in these GBM cells identified by Neftel et al. Another study revealed that aneuploid cells were found in only 68% and 32% of samples of the C and P regions, respectively, and they comprised 3% to 84% of all cells (4). Recurrent GBM cells develop novel genetic variations compared to primary GBM cells under the selective pressure of radiotherapy and chemotherapy (7,8,49,50). However, further research is needed to elucidate the dynamic changes in CNV occurrence and progression, as well as the underlying regulatory mechanism.

Although the generation of CNVs is a pivotal mechanism used by cancer cells to regulate gene expression, the transcriptome is orchestrated by multiple factors. Only 40-60% of the amplified genes in cancer cells correspond to RNA transcripts (51). Therefore, our inferred CNVs based on scRNA-seq contained information on both CNVs and other regulatory mechanisms of the transcriptome and omitted some CNVs inconsistent with the corresponding RNA transcript. Our data revealed that GBM cells of different CNV patterns were also controlled by different transcriptional regulatory networks. Despite prominent heterogeneity, we identified some common TFs among the four CNV patterns, such as JUN and MYC, which were also identified as central TFs by a recently published study using multiomics to construct a kinase-TF centered network in HGG (52). These common TFs might have essential roles in GBM cells. Notably, the core TF–TCF gene set of CNV pattern 4 was negatively correlated with both the OS and PFS of patients with GBM, indicative of CNV pattern 4 GBM cell as a promising prognostic marker and therapeutic target.

Bulk transcriptomic profile analyses have classified GBM cells into three molecular subtypes: the classical, mesenchymal, and proneural subtypes (8,11). To simplify this model, Wang et al. focused on the glioma stem-like cells (GSCs) of the mesenchymal, proneural, and classical subtypes (mGSC, pGSC, and cGSC, respectively) (53), and found that the cellular states of GSCs ranged from mesenchymal root to proneural terminal, between which was an intermediate population. Interestingly, three TFs specific to this intermediate population, ATF3, JUNB, and JUN, were also found in the core TF–TCF gene set of CNV pattern 4. This consistency further supports our hypothesis that CNV pattern 4 might be a shared transitional state of heterogeneous GBM cells. We also learned from published studies that ATF3 is a bidirectional switch that regulates the treatment responses of GBM cells. On the one hand, ATF3 is involved in resistance to temozolomide (TMZ), cisplatin, and ultraviolet light (54,55), but on the other hand, it serves as a key mediator in proteasome inhibitor-induced GSC-selective apoptosis (56). In addition, another two CNV pattern 4-specific core TFs, FOS and JUNB, have been shown to confer GBM cells with TMZ resistance (57). These reports underscore the intrinsic resistance of CNV pattern 4 GBM cells to chemotherapy, as well as the necessity of more research to overcome this obstacle.

Remarkably, a sub-cluster of CNV pattern 4 GBM cells expressed CD133 and showed the ability to form spheres, indicating stem cell-like properties and tumorigenic potential. Consistent with these results, Piccirillo et al. showed that GBM cells isolated from the P/N regions could form neoplasms in a mouse model, although they had lower expression levels of the stem cell markers Nestin, SOX2, and Notch2 compared to those in the C region (25). In addition, CD133-positive and Nestin-positive invasive GBM cells have been detected in murine models and shown to exhibit self-renewal ability (58,59). CD133-positive GBM stem cells (GSCs) are the source of VM (41,42) that leads to a compromised effect of antiangiogenesis therapy in GBM (60). In this study, we found that a sub-cluster of CNV pattern 4 GBM cells resembled (to varying degrees) endothelial cells and pericytes, which implies the possibility of CNV pattern 4 GBM cells participating in VM. Given that VM might contribute to the resistance to antiangiogenesis therapy, this sub-cluster of CNV pattern 4 GBM cells would be an ideal target of postoperative treatment.

Considering the infiltrative property of GBM cells and the disruption of BBB in GBM lesion, we also examined PBMCs from patients with GBM to explore whether GBM cells exist in circulation. We discovered a subset of proliferative cells that were initially regarded as malignant cells but were finally demonstrated to be lymphocytes. These proliferative lymphocytes were found in the PBMCs from patients with GBM but not in healthy volunteers or patients with epilepsy. In addition to being refractory to standard care, GBM is notorious for being highly immunosuppressive both locally and systemically (61,62), which makes immunotherapy difficult in GBM. These proliferative lymphocytes, although comprising <1% of PBMCs, might represent a nascent immune response to GBM and could help develop a new way to identify the potential of the immune system in GBM treatment. Besides, because these proliferative lymphocytes are specific to the PBMCs from patients with GBM, they could be suitable targets for monitoring recurrence in addition to MRI, which has limited efficiency in differentiating tumor recurrence and pseudoprogression (63). Therefore, the biological functions and mechanisms of these proliferative immune cells warrants future, detailed exploration.

In conclusion, this study has identified an unusual GBM cell subset that is predominantly found in the P and N regions and is genomically distinct from GBM cells in the C region. These cells have a fragmented, atypical CNV profile. However, these cells resemble GBM cells with typical CNVs at the transcriptomic level to a certain extent, and have the potential to transform into their respective cellular states. A sub-cluster of these cells possesses stem cell-like properties and might, therefore, participate in VM. Finally, yet importantly, the core signature of the transcriptional regulatory network of these cells negatively correlates with GBM patient outcomes. In parallel, we discovered proliferative immune cells that were highly responsive to GBM in both GBM tissue and PBMCs. The unusual CNV 4 GBM cells and proliferative immune cells from patients with GBM identified in this study might be ideal targets for GBM diagnosis, therapy and prognosis.

## METHODS

### Sample acquisition

This study was approved by the Ethics Committee of Nanfang Hospital. Written informed consent was provided by all patients. Peripheral blood samples were obtained before surgery at Nanfang Hospital, Southern Medical University. GBM samples were obtained under MRI guidance during surgery. Normal brain samples were obtained during surgical treatment for ependymoma. The diagnosis of IDH1-wt GBM was confirmed by the Department of Pathology of Nanfang Hospital (Supplementary Table S1: Pathological diagnosis information of four patients with GBM).

### HE and IHC staining

HE and IHC staining assays were performed as previously reported (64,65). Briefly, specimen slices were cut from paraffin-embedded tissue blocks with a microtome (Leica, EG1150H, Wetlzar, Germany), deparaffinized and rehydrated. For HE staining, the slices were sequentially immersed in HE. For IHC staining, the slices were further processed, including antigen retrieval, the blocking of endogenous peroxidase, primary antibody incubation, secondary antibody incubation, and nuclear staining. Finally, the slices were sealed with mounting medium for imaging. The antibodies used in this study are listed in Supplementary Table S5.

### Primary GBM cell culture and immunocytochemistry (ICC)

GBM samples were maintained on ice and sent to the laboratory within 1 h of isolation. Samples were minced with a scalpel and then enzymatically digested with a Tumor Dissociation Kit (Miltenyi Biotec, 130-095-929, Germany) on a gentleMACS Octo Dissociator with Heaters (Miltenyi Biotec, 130-096-427, Germany) according to the manufacturer’s instructions. To isolate stem-like GBM cells, primary GBM cells were cultured in DMEM/F-12 medium (Thermo Fisher Scientific, Gibco, #8117165, Shanghai, China) supplemented with B27 (1×, Thermo Fisher Scientific, #17504044, Shanghai, China), EGF (20 ng/ml, Promokine, #C-60170, Guangzhou Jetway Biotechnology Company, Guangzhou, China), and FGF (20 ng/ml, Promokine, #C-60240, Guangzhou Jetway Biotechnology Company, Guangzhou, China), in which some GBM cells without stem cell-like properties would be degraded. ICC was performed as previously reported (64).

### Sample preparation and 10× Genomics scRNA-seq

GBM tissue samples were washed twice with pre-cooled Hank’s balanced salt solution (HBSS, Thermo Fisher, 88284, USA), during which apparent vessels and pia matter were removed. Clean samples were dissociated mechanistically and enzymatically, as described above (Miltenyi Biotec, 130-095-929; gentleMACS Octo Dissociator with Heaters, 130-096-427, Germany). Peripheral blood samples from patients with GBM, patients with epilepsy, and healthy donors were collected into EDTA tubes. PBMCs were isolated with Ficoll-Plaque Premium (GE Healthcare, 17544203-1, USA) according to the manufacturer’s protocol. First, 4 mL peripheral blood was slowly added to 3 mL Ficoll solution, which was then centrifuged at 400×g for 30 min at 18°C. Next, the PBMC layer was transferred to another tube, washed twice with PBS and centrifuged at 400×g for 15 min at 18°C.

Subsequently, cells from GBM tissue samples or peripheral blood samples were resuspended in DMEM (Thermo Fisher, 11320033, USA) containing 10% fetal bovine serum (FBS) (Thermo Fisher, 10099141, USA) and filtered through a 70-μm nylon filter (Corning Falcon, 431751, USA). The filtered cells were subjected to 1× Red Blood Cell Removal Solution (Biogems, 64010-00-100, USA) for 5 min to remove red blood cells, and then the cell debris was removed with Debris Removal Solution (Miltenyi, 130-109-398, Germany). Following two washes and resuspension in PBS solution containing 0.04% bull serum albumin (BSA) (Thermo Fisher, AM2616, USA), the cells were mixed with Trypan Blue (Thermo Fisher, T10282, USA) to assess their viability using a hemocytometer (Thermo Fisher, C10312, USA). Then, the appropriate volume for each sample was calculated for a target capture of 6,000, cells according to the user guide of the Single Cell 3’ Reagent Kit v2 (10× Genomics company, 120237-16, USA). Single-cell droplet generation, reverse transcription, and cDNA library preparation were performed according to the manufacturer’s protocols. Finally, the libraries were sequenced on an Illumina NovaSeq 6000 with 150 bp paired-end sequencing. A median sequencing depth of 50,000 reads/cell was targeted for each sample.

### scRNA-seq data processing and the identification of non-malignant cells

Raw sequencing data (bcl files) were converted to fastq files with Illumina bcl2fastq, version 2.19.1 and aligned to the human genome reference sequence (http://cf.10×genomics.com/supp/cell-exp/refdata-cellranger-GRCh38-1.2.0.tar.gz). The CellRanger 2.2.0 (10× Genomics) analysis pipeline was used to generate a digital gene expression matrix from these data according to its guidelines. The raw digital gene expression matrix (UMI counts per gene per cell) was filtered, normalized, and clustered using R 3.5.2 software (https://www.R-project.org/). Cell and gene filtering was performed as follows. For each detected cell, UMIs were less than the (1-doublet rate) to exclude multiplets. The multiplet rates were determined according to the user guide provided by 10× Genomics. The UMI was larger than the 8th percentile to exclude ambient bias (66). Mitochondrial RNA was less than the 90th percentile to exclude devitalized cells. For each sample, the number of detected cells was controlled within 6,000 by adjusting the baseline gene number. Then, normalization and centralization were performed according to the detected genes in each cell, and outlier cells with z-scores exceeding ±6 were excluded. We utilized the “NormalizeData” and “ScaleData” functions in the Seurat package to normalize and scale the single-cell gene expression data. The highly variable genes (HVGs) across the single cells in each sample were determined using the Seurat “FindVariableGenes” function. Canonical correlation analysis (CCA) was applied to correct for batch effects observed in the twelve GBM tissue samples and the four PBMC samples (67). For CCA, the union gene sets from the top 2,000 HVGs of each sample was used, and the combined data were aligned using the first 30 CC dimensions. Aligned cells were clustered with the Seurat “FindClusters” function using the first 20 aligned CC dimensions at a resolution of 0.5. Clustering results were visualized using t-distributed stochastic neighbor embedding (t-SNE). The DEGs of each cluster were identified by comparison with the remainder of the other clusters. Cell type-specific markers and GO enrichment results were combined to classify the identity of each cluster. Using these approaches, oligodendrocytes, macrophages, T cells and putative GBM cells were identified in GBM samples. T cells, B cells, and monocytes were identified in PBMC samples. Data analysis was performed using standard packages, including Seurat 2.3.4 and Monocle 2.6.4, as well as website platforms such as GO: http://geneontology.org/) (68,69), Metascape (http://metascape.org/gp/index.html#/main/step1) (70), and Gene Expression Profiling Interactive Analysis (GEPIA2: http://gepia2.cancer-pku.cn/#index) (71).

### CNV inference from scRNA-seq data

The CNVs of putative GBM cells were evaluated according to transcriptomic information by the moving average algorithm inferCNV (6,10,27,28,72,73). Briefly, all detected genes were sorted by their chromosomal location, and then the expression level of each gene in each cell was calculated based on an additional 100 neighboring genes (50 upstream and 50 downstream); this parameter was set as the sliding window for each chromosome. In this way, gene-specific expression patterns could be smoothed to a certain extent, and the effects of the CNVs on the transcriptome became prominent. Considering that the cell lineages of GBM belong to the central nervous system, only oligodendrocytes were used as the normal karyotype reference. Hierarchical clustering was performed by “fastcluster 1.1.25” to identify CNV patterns (74).

### Construction of the TF-TCF-target regulatory network

Gene lists of the TFs and TCFs were obtained from the Animal Transcription Factors Database (http://bioinfo.life.hust.edu.cn/AnimalTFDB/#!/) (75–77). Then, the gene lists of the TFs and TCFs were intersected with the DEGs of the four CNV patterns to identify CNV pattern-specific TFs and TCFs. Interactions between the selected TFs and TCFs were identified with the STRING database (https://string-db.org/) (78). To identify a solid interaction relationship, only interactions from experiments and databases with a minimum required interaction score of 0.9 were considered. TF–TCF interaction networks were constructed with Cytoscape software based on these confident interactions. Then, the interaction degree of each TF was calculated to identify the hub TFs for each CNV pattern. The top three TFs with the highest degrees of interaction were chosen for target gene analysis in Cistrome Data Browser (http://cistrome.org/db/#/) (79,80). The target gene lists were intersected with the DEGs of the four CNV patterns to select the differentially expressed target genes (DETGs). Finally, the hub TFs, their neighbors in the TF–TCF interaction network and the DETGs were used to construct a simplified TF–TCF-target regulatory network for each CNV pattern in Cytoscape software.

### Data availability

The scRNA-seq data described in this study are available upon reasonable request from the corresponding authors.

### Statistics

The statistical analyses were performed in SPSS statistical software, version 21.0 (SPSS, Inc., Chicago, IL, USA). Survival analysis of the data from the TCGA GBM cohort was performed in the GEPIA2 platform, which used the log-rank test for the hypothesis test (71). A rank-sum test was performed for RRG comparisons between primary GBM and recurrent GBM in the China Glioma Genome Atlas (CGGA) database (http://www.cgga.org.cn/index.jsp) (Figure S13C). The Spearman correlation coefficient of the RRGs was calculated using the GBM cohort (190 patients) from CGGA (Figure S13D). Significance cut-off: *, P <0.05; **, P < 0.01; ***, P < 0.001; and ****, P < 0.0001.

## Funding

This study was supported by the National Natural Science Foundation of China (Grant nos. 81773290, 81802505), Key R&D Program of Guangdong Province (2018B090906001), Guangdong Science and Technology Department (2017A030313497), PLA Logistics Research Project of China (18CXZ030, CWH17L020), Science and Technology Planning Project of Guangzhou (201902020017).

## Acknowledgements

We are grateful to Peng Fang, Mei Zheng, Jiaohua Jiang and their colleagues of Guangzhou SALIAI Stemcell Science and Technology Co., Ltd. (Guangzhou, China) for helpful discussion about this study with them.

## Contributions

Conception and design of the work: YWL, STQ, RWY, and GLH.

Acquisition, analysis and interpretation of data: RWY, GLH, JLG, YML, HMS, KSL, GZY, ZYL, XRW, SDX, ZFL, XFS, KL, STQ, YWL.

Drafting of the manuscript: RWY and YWL.

Critical revision for important intellectual content: RWY, GLH, JLG, YML, STQ, YWL.

All authors approved the final version of this manuscript.

## Supplementary figure legends

**Figure S1. A,** HE staining and IHC of 12 samples from four patients with GBM. The C region was characterized by the histological hallmarks of GBM, including microvasculature proliferation and necrosis, as well as a high ratio of KI67-positive cells. The P and N regions resembled normal brain tissue to varying extents. **B,** HE and IHC stains of normal brain tissue obtained during the surgical approach to ependymoma.

**Figure S2. A,** The CNV pattern compositions of the different transcriptomic clusters. **B,** The connections between CNV patterns and transcriptomic clusters. The widths of the connecting lines correspond to the number of cells. **C,** The developmental trajectory of GBM cells inferred by the Monocle2 algorithm. The distributions of different groups along three state axes are shown separately. **D,** The top six DEGs among the three states. **E,** The pathway enrichment results of CNV pattern 1-specific DEGs. **F,** The pathway enrichment results of CNV pattern 4-specific DEGs. The pathways involved in the regulation of gene expression are marked in red rectangles.

**Figures S3.** TF–TCF interaction networks of the four CNV patterns. TFs are listed in circles, and TCFs are listed in rounded rectangles. The sizes of the TFs and TCFs were determined by the number of their connecting neighbors, which was defined as the interaction degree. Upregulated TFs and TCFs are indicated in red, while downregulated TFs and TCFs are indicated in blue. A larger absolute FC value corresponds to a deeper color. The core gene sets of the TF-TCF interaction networks are listed on the right side, and the parameters are listed on the bottom.

**Figures S4.** TF–TCF interaction networks of the four CNV patterns. TFs are listed in circles, and TCFs are listed in rounded rectangles. The sizes of the TFs and TCFs were determined by the number of their connecting neighbors, which was defined as the interaction degree. Upregulated TFs and TCFs are indicated in red, while downregulated TFs and TCFs are indicated in blue. A larger absolute FC value corresponds to a deeper color. The core gene sets of the TF-TCF interaction networks are listed on the right side, and the parameters are listed on the bottom.

**Figure S5.** TF–TCF interaction networks of the four CNV patterns. TFs are listed in circles, and TCFs are listed in rounded rectangles. The sizes of the TFs and TCFs were determined by the number of their connecting neighbors, which was defined as the interaction degree. Upregulated TFs and TCFs are indicated in red, while downregulated TFs and TCFs are indicated in blue. A larger absolute FC value corresponds to a deeper color. The core gene sets of the TF-TCF interaction networks are listed on the right side, and the parameters are listed on the bottom.

**Figure S6.** TF–TCF interaction networks of the four CNV patterns. TFs are listed in circles, and TCFs are listed in rounded rectangles. The sizes of the TFs and TCFs were determined by the number of their connecting neighbors, which was defined as the interaction degree. Upregulated TFs and TCFs are indicated in red, while downregulated TFs and TCFs are indicated in blue. A larger absolute FC value corresponds to a deeper color. The core gene sets of the TF-TCF interaction networks are listed on the right side, and the parameters are listed on the bottom.

**Figures S7.** Simplified TF–TCF-target regulatory networks of the four CNV patterns. TFs are listed in the inner layer (circles), TCFs are listed in the second layer (diamond), and TF targets are listed in the outermost layer (rounded rectangles). Upregulated expression is indicated in red, while downregulated expression is indicated in blue. A larger absolute FC value corresponds to a deeper color. Interactions between TFs and TCFs are depicted by straight lines. TFs and their corresponding targets are connected by lines, with a single arrow pointing to the target.

**Figures S8.** Simplified TF–TCF-target regulatory networks of the four CNV patterns. TFs are listed in the inner layer (circles), TCFs are listed in the second layer (diamond), and TF targets are listed in the outermost layer (rounded rectangles). Upregulated expression is indicated in red, while downregulated expression is indicated in blue. A larger absolute FC value corresponds to a deeper color. Interactions between TFs and TCFs are depicted by straight lines. TFs and their corresponding targets are connected by lines, with a single arrow pointing to the target.

**Figures S9.** Simplified TF–TCF-target regulatory networks of the four CNV patterns. TFs are listed in the inner layer (circles), TCFs are listed in the second layer (diamond), and TF targets are listed in the outermost layer (rounded rectangles). Upregulated expression is indicated in red, while downregulated expression is indicated in blue. A larger absolute FC value corresponds to a deeper color. Interactions between TFs and TCFs are depicted by straight lines. TFs and their corresponding targets are connected by lines, with a single arrow pointing to the target.

**Figures S10.** Simplified TF–TCF-target regulatory networks of the four CNV patterns. TFs are listed in the inner layer (circles), TCFs are listed in the second layer (diamond), and TF targets are listed in the outermost layer (rounded rectangles). Upregulated expression is indicated in red, while downregulated expression is indicated in blue. A larger absolute FC value corresponds to a deeper color. Interactions between TFs and TCFs are depicted by straight lines. TFs and their corresponding targets are connected by lines, with a single arrow pointing to the target.

**Figure S11. A,** The core gene sets of the TF–TCF interaction networks of the four CNV patterns. The color gradation shows the interaction degrees of the DEGs. Common TFs and TCFs in different CNV patterns and CNV pattern-specific TFs/TCFs are arranged from left to right. **B,** An expression analysis of CNV pattern-specific TFs based on the TCGA and GTEx databases. **C-H,** Expression analysis of CNV pattern-specific TFs among the four molecular subtypes of GBM samples in the TCGA database.

**Figure S12. A-D,** Expression analysis of CNV pattern 4-specific TFs, JUNB and FOS, in different types of malignancies included in the TCGA database. GBM, glioblastoma. LAML, acute myelocytic leukemia. PAAD, pancreatic adenocarcinoma. BLCA, bladder urothelial carcinoma. DLBC, lymphoid neoplasm diffuse large B-cell lymphoma. LUAD, lung adenocarcinoma. LUSC, lung squamous cell carcinoma. OV, ovarian serous cystadenocarcinoma. THYM, thymoma. LGG, low-grade glioma. ACC, adrenocortical carcinoma. BRCA, breast invasive carcinoma. HNSC, head and neck squamous cell carcinoma. KICH, kidney chromophobe. KIRP, kidney renal papillary cell carcinoma. LIHC, liver hepatocellular carcinoma. SKCM, skin cutaneous melanoma.

**Figure S13. A and B,** The prognostic value of the CNV pattern core gene sets on the OS **(A)** and DFS **(B)** of patients with GBM. **C,** The differential expression of CNV pattern 4-specific genes JUNB, ATF3, ACTN1, CEBPB, EGR1, and NFKBIA in primary and recurrent GBM samples included in the China Glioma Genome Atlas (CGGA) database. **D,** Spearman’s correlation analysis of RRGs of CNV pattern 4 based on the GBM cohort from CGGA database.

**Figure S14. A,** The CNV pattern compositions of different samples from GBM-4 patients. **B,** Pathway enrichment results of the DEGs of sub-cluster 3. **C,** Three-dimensional coordinate diagram of the endothelial cell score (X-axis), pericyte score (Y-axis), and stemness score (Z-axis). GBM cells of CNV pattern 4 were scored in these three dimensions and then plotted in the coordinate frame. **D,** Developmental trajectory of CNV pattern 4 GBM cells inferred by the Monocle2 algorithm. Distributions of sub-clusters along the three state axes are presented separately.

**Figure S15. A,** IHC staining of SOX2, Nestin, and GFAP in GBM samples from the C, P, and N regions. **B,** IHC staining of SOX2, Nestin, and GFAP in normal brain tissue obtained during the surgical approach to ependymoma.

**Figure S16. A,** A t-SNE plot of 23,850 immune cells isolated from 12 GBM tissue samples. **B,** GO enrichment analysis of intracranial immune cell sub-cluster 17 DEGs compared to GBM cells. **C-E,** The expression of markers of proliferative cells, malignant cells, and monocytes in the PBMCs of patients with GBM. **F,** GO enrichment analysis of PBMC sub-cluster 10 DEGs compared to GBM cells. **G and H,** The expression of proliferation-associated genes in the PBMCs of healthy volunteers and patients with epilepsy.

## Notes

**Competing interests:** The authors declare no potential conflicts of interest.

#### Summary of Updates

Figure 5 and Figure S16, as well as the manuscript have been revised.

