## Supplementary figures and images for "Genetic characterization of stem cell-like cancer cell in the peri-tumoral regions and proliferative lymphocyte in the peripheral blood of patients with glioblastoma"

### Supplementary Figure S1

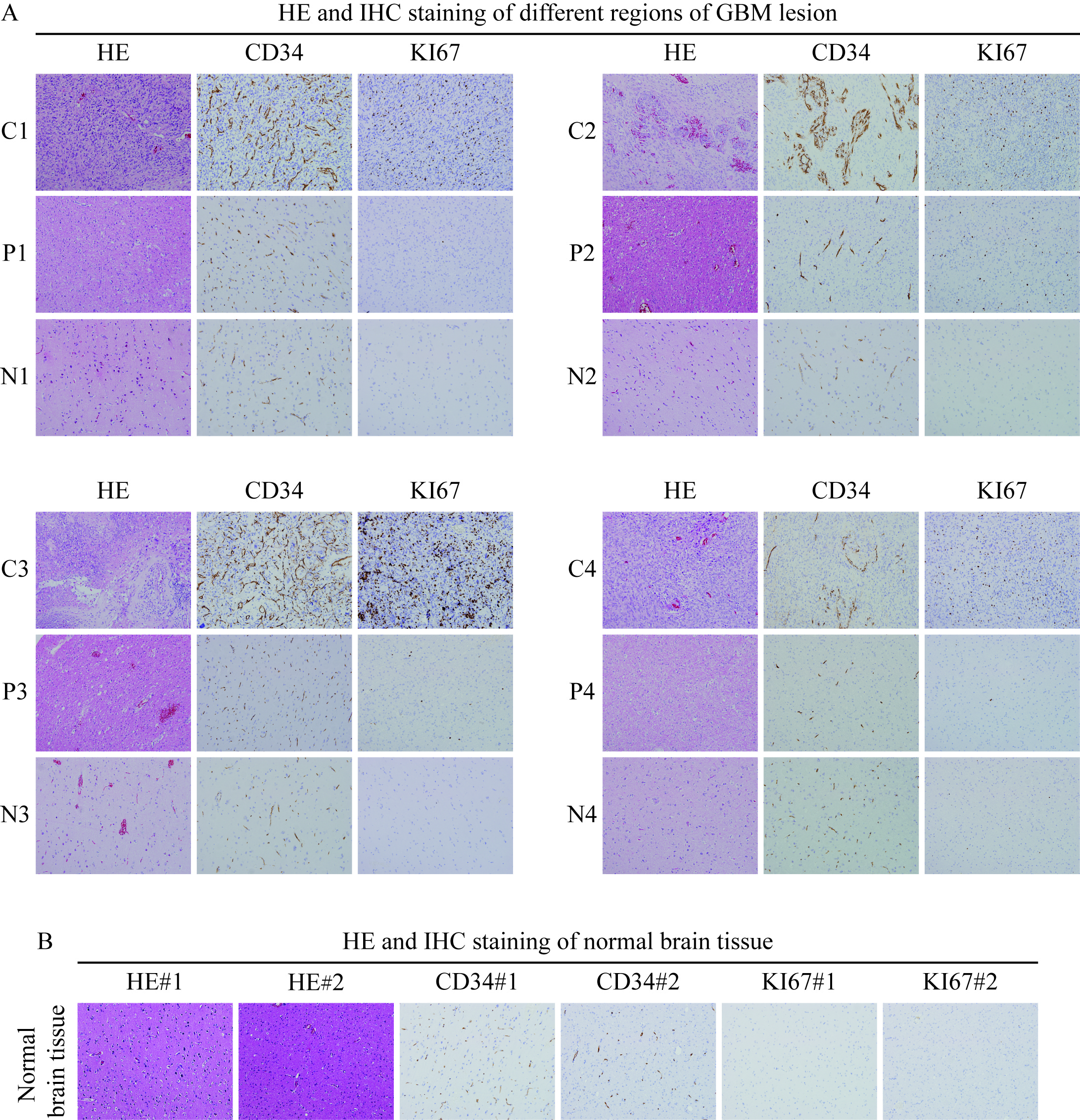

### Supplementary Figure S2

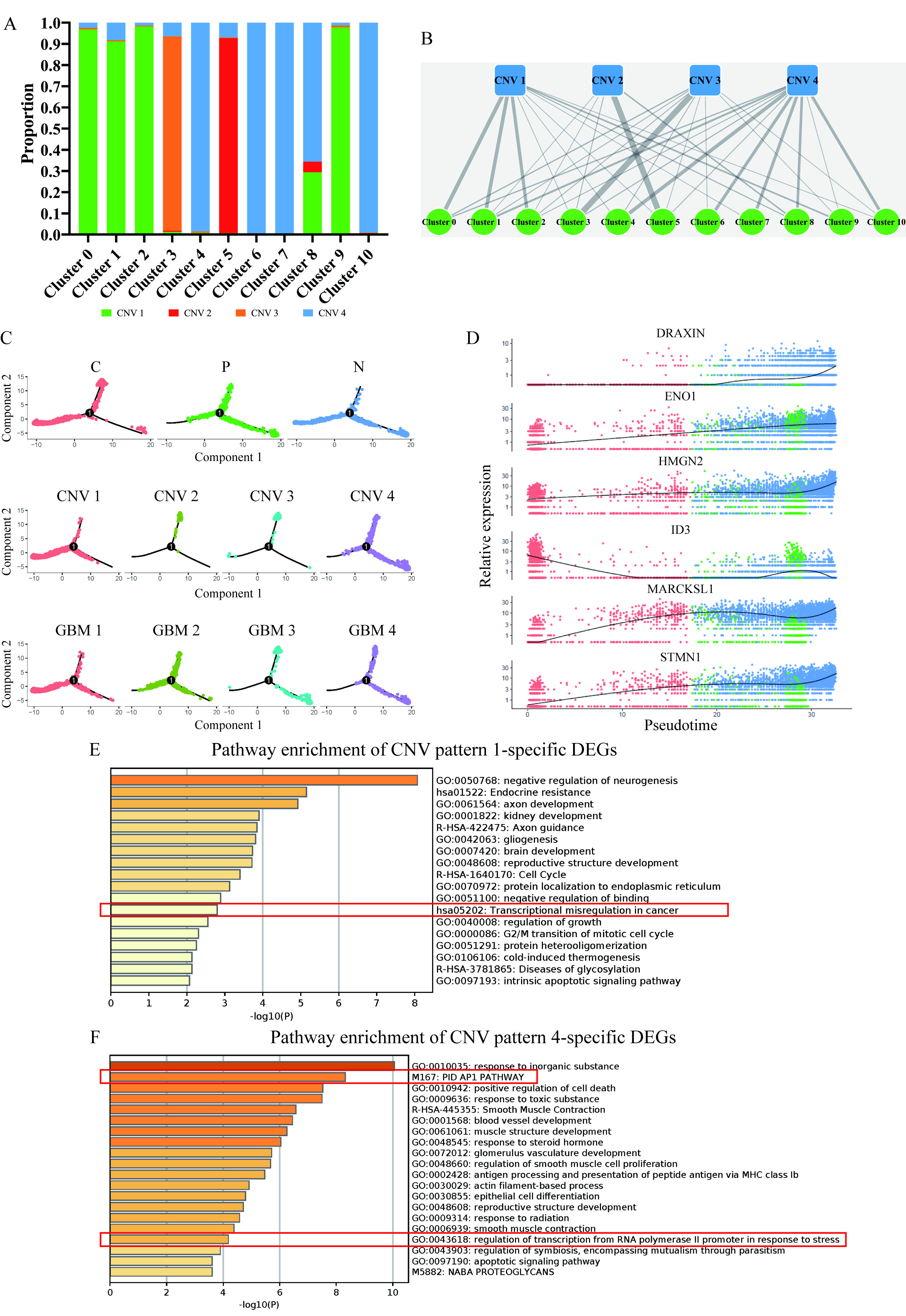

### Supplementary Figure S3

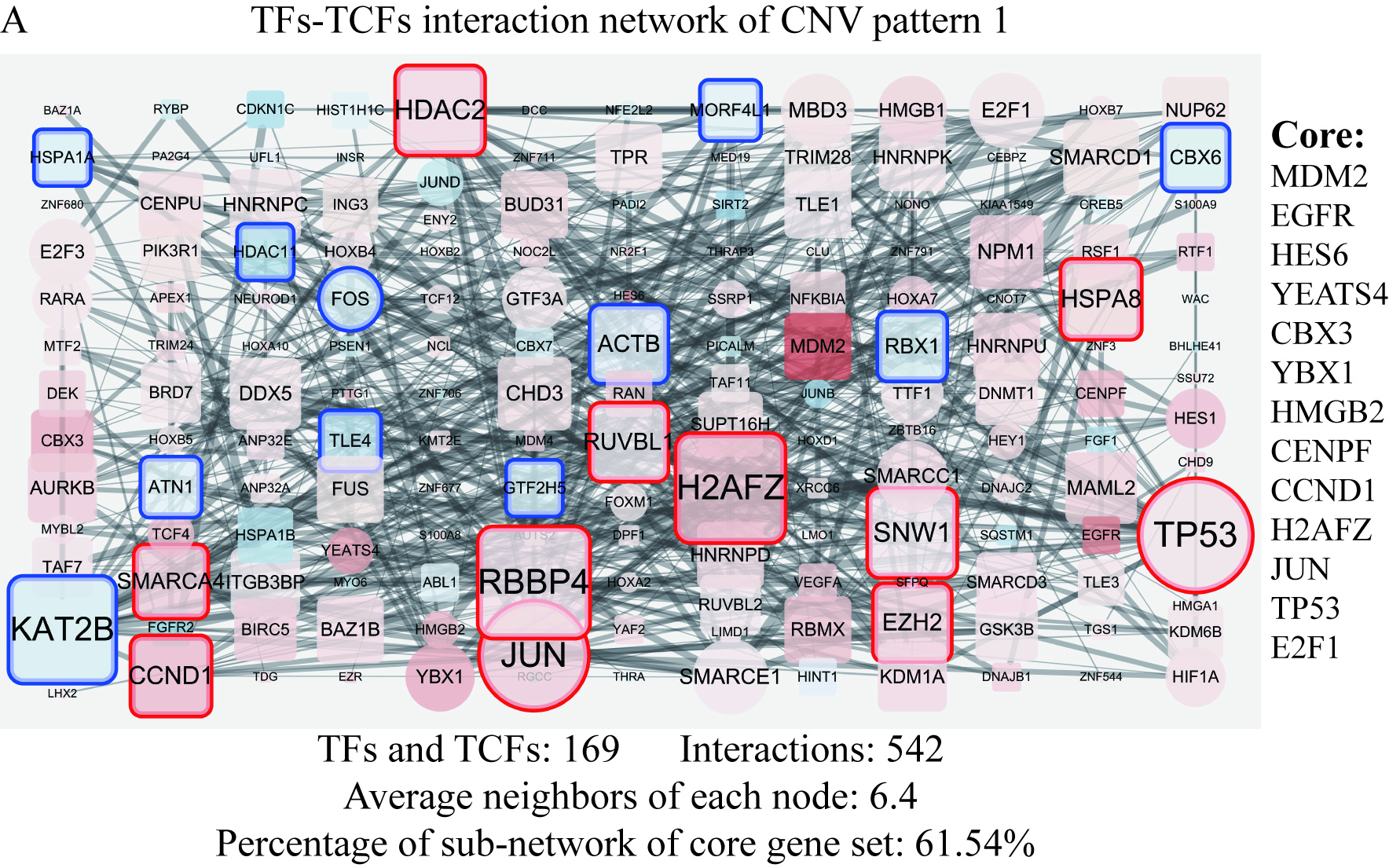

### Supplementary Figure S4

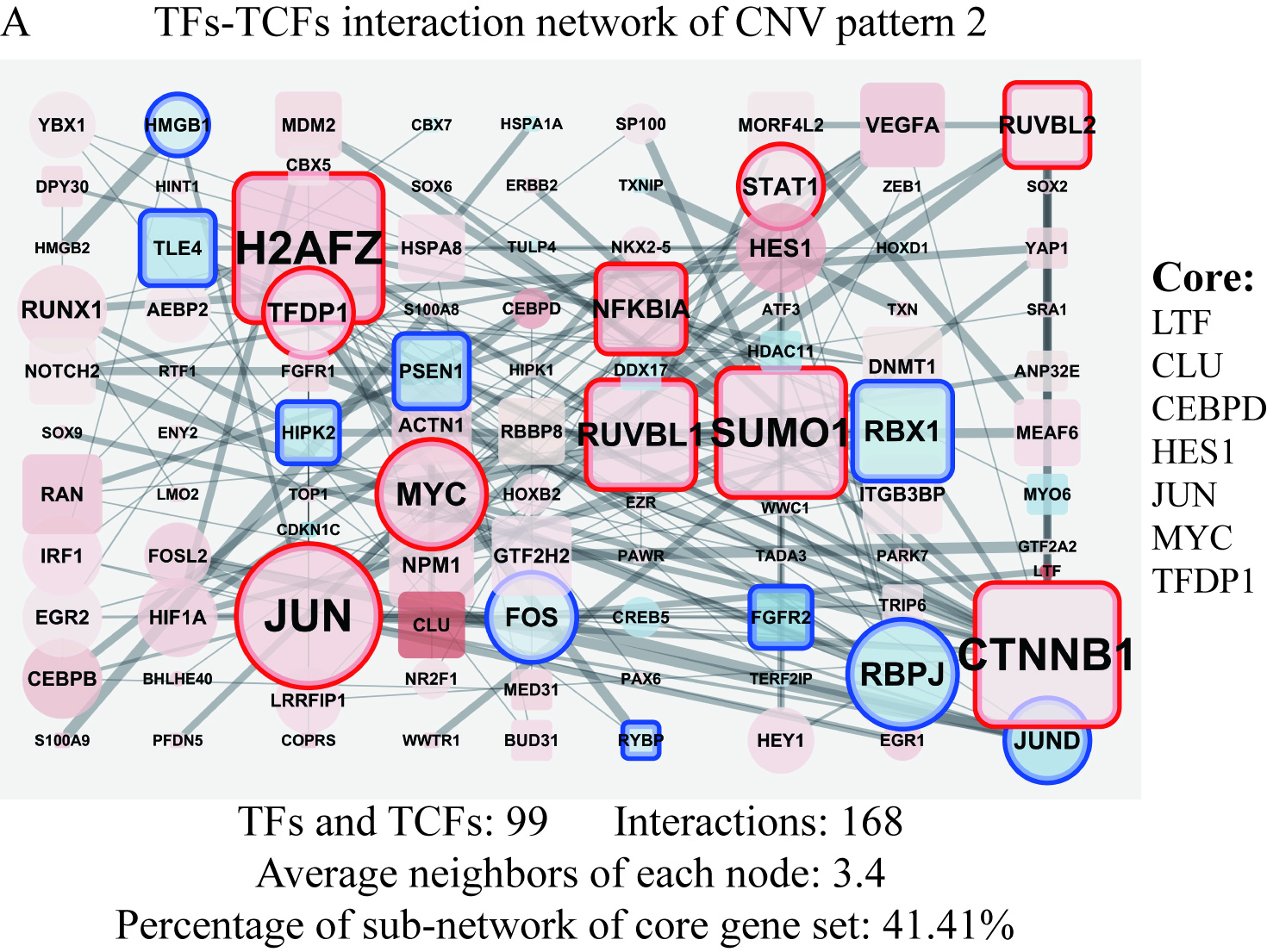

### Supplementary Figure S5

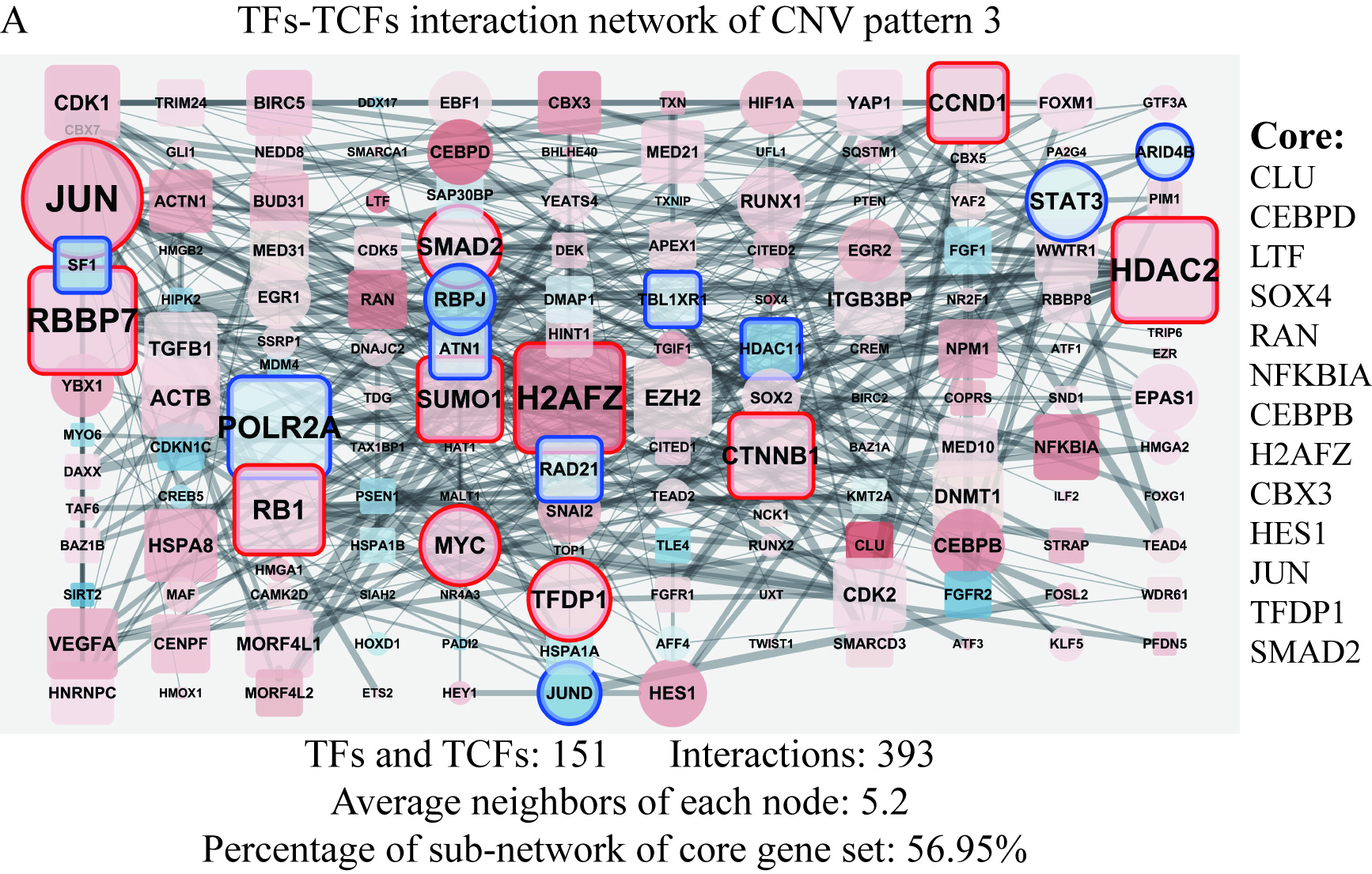

### Supplementary Figure S6

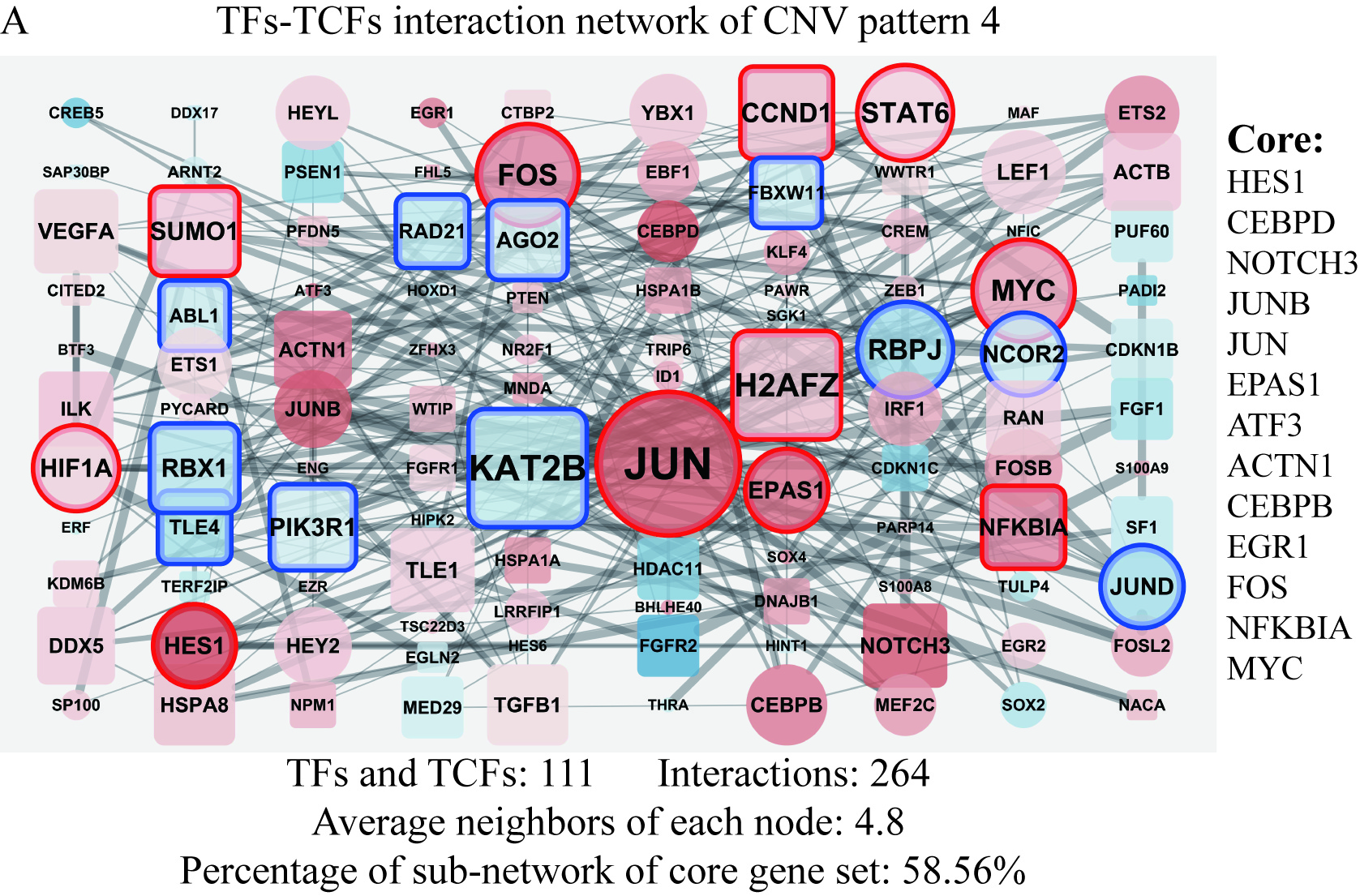

### Supplementary Figure S7

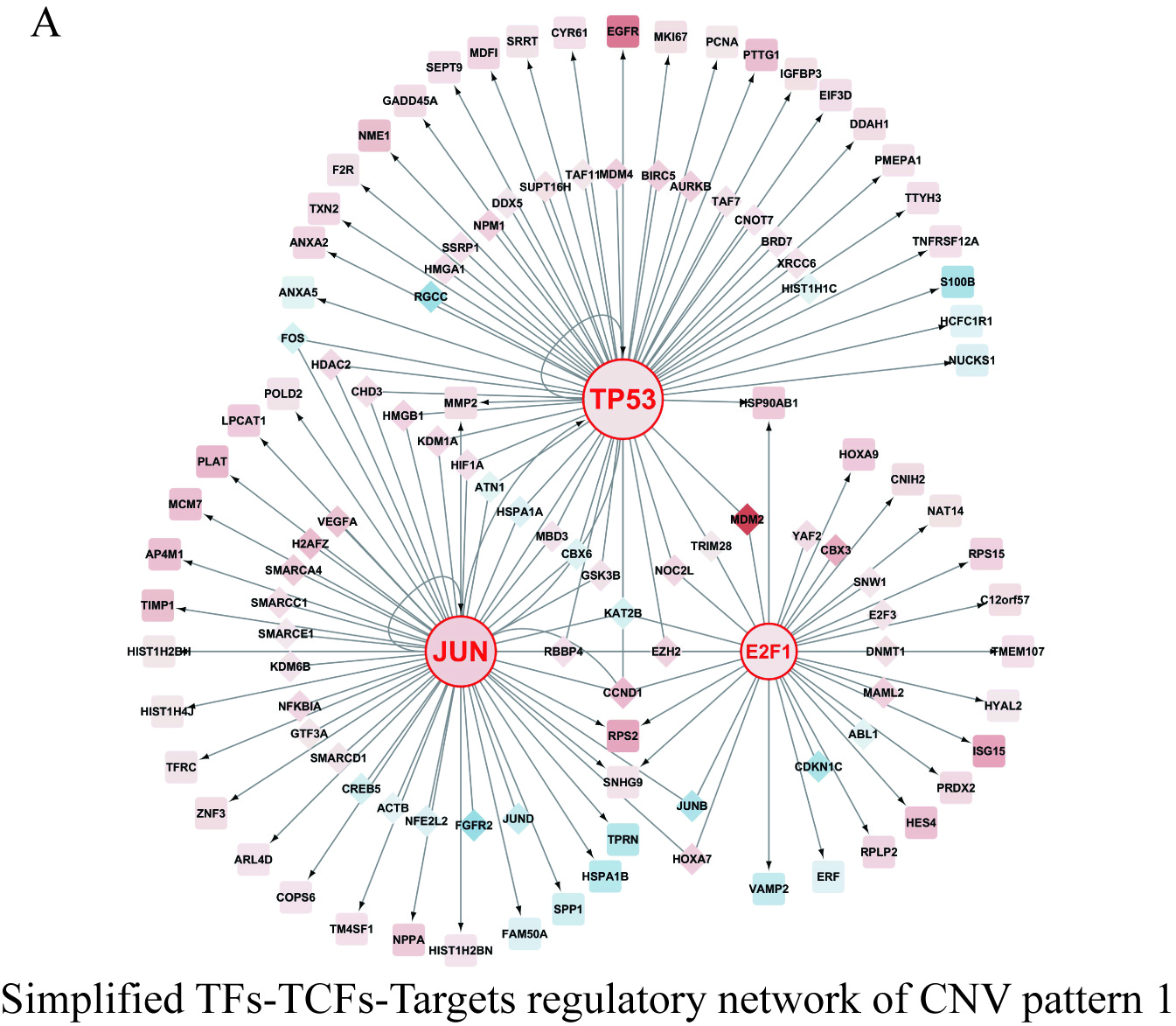

### Supplementary Figure S8

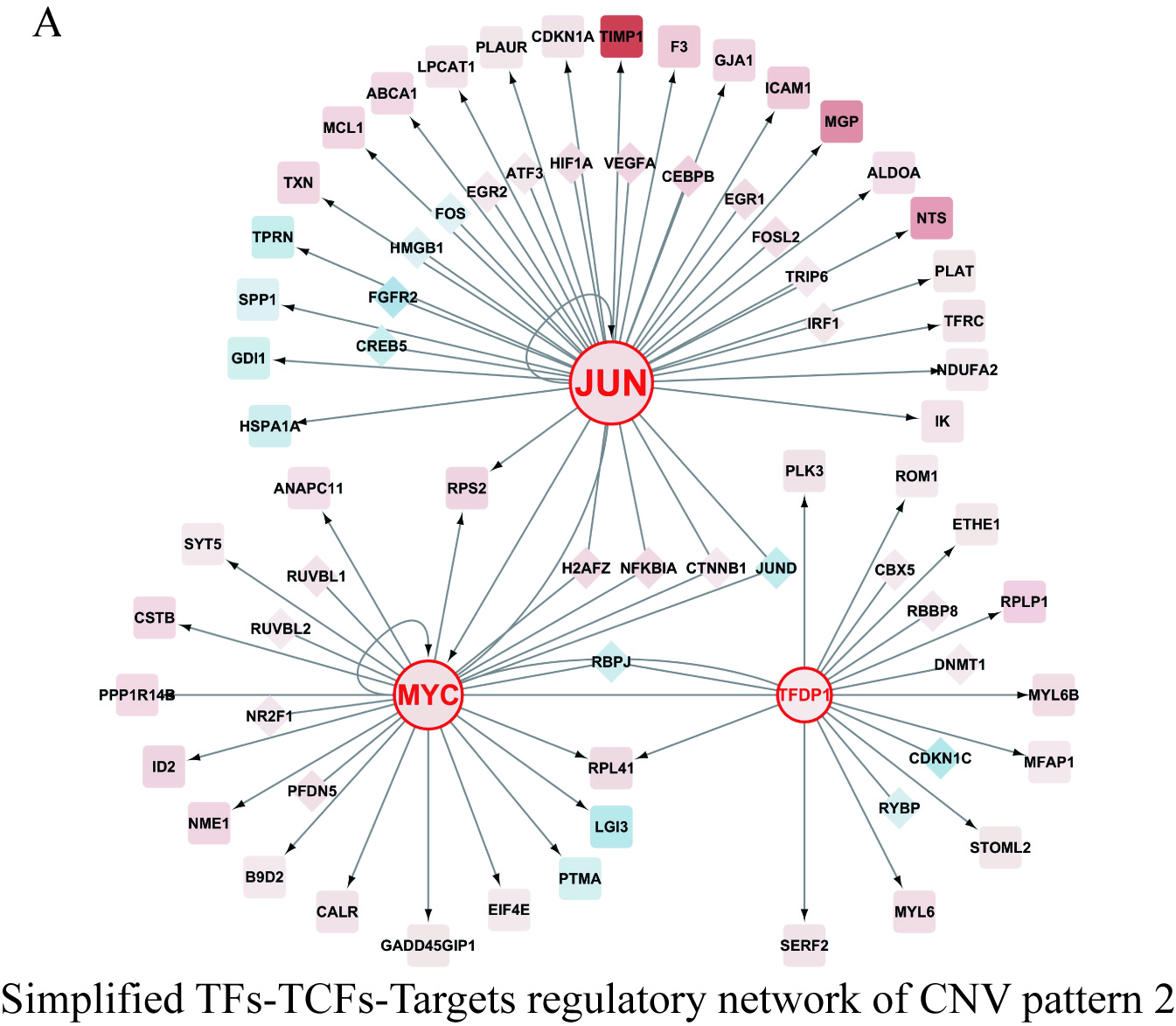

### Supplementary Figure S9

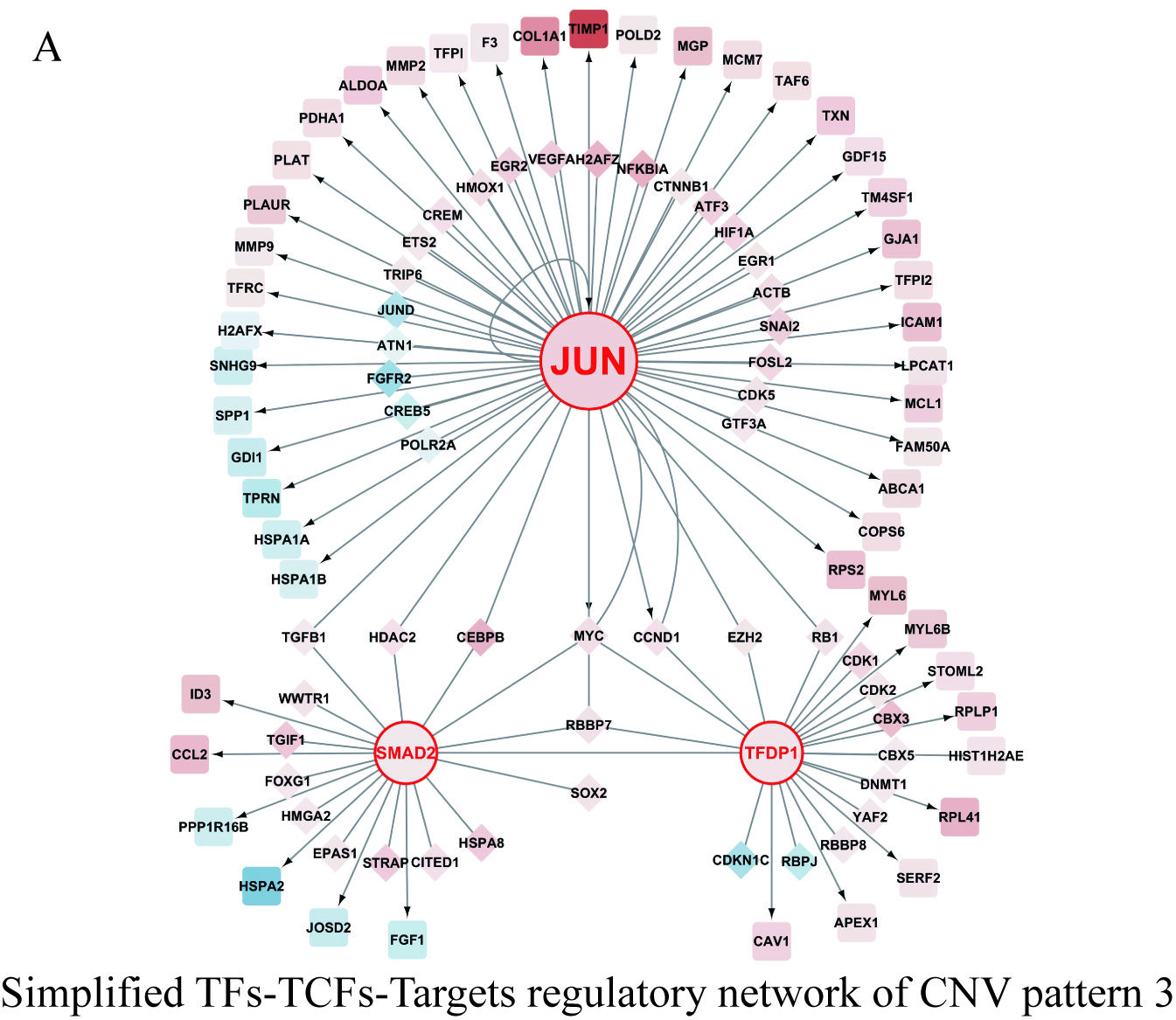

### Supplementary Figure S10

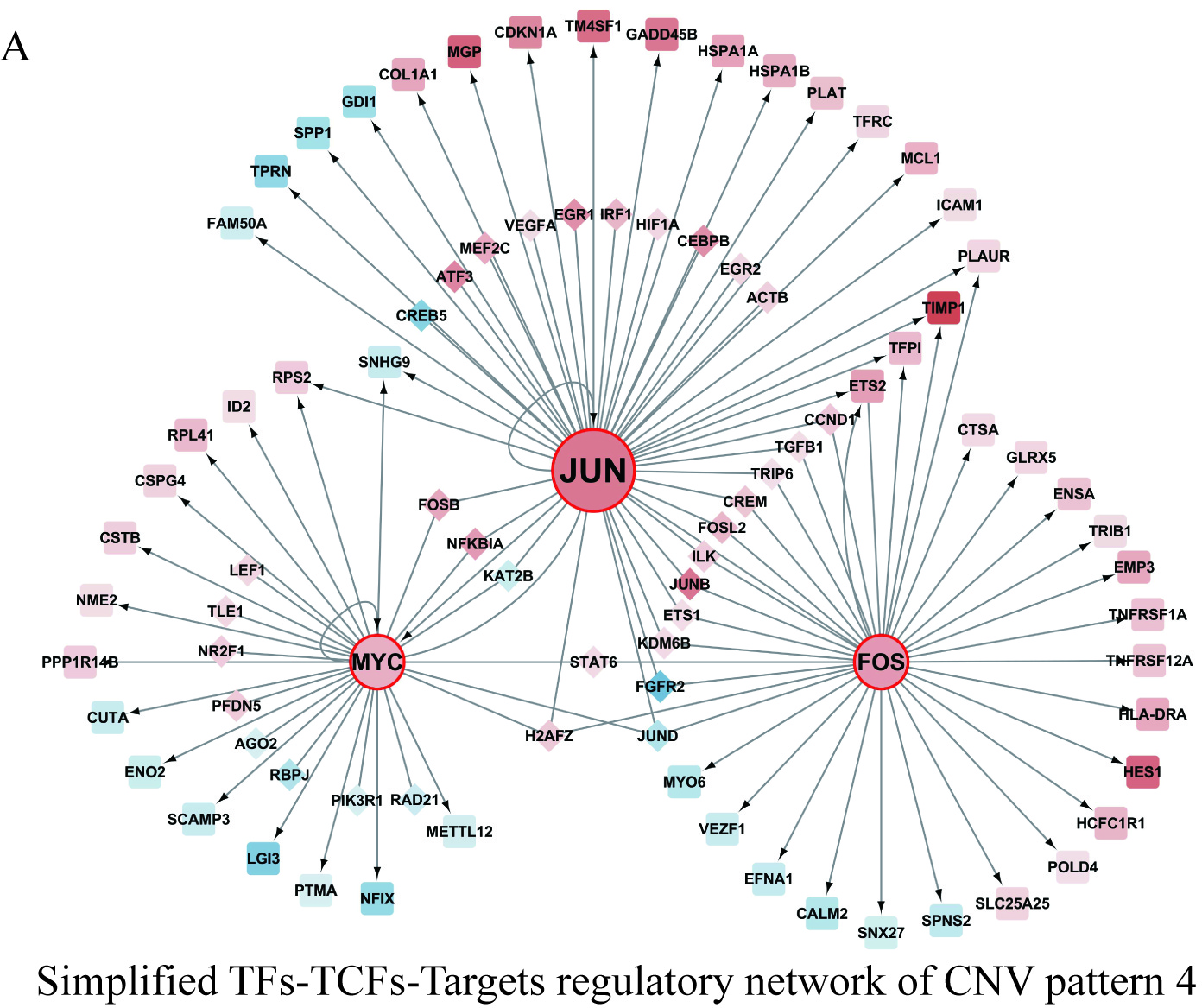

### Supplementary Figure S11

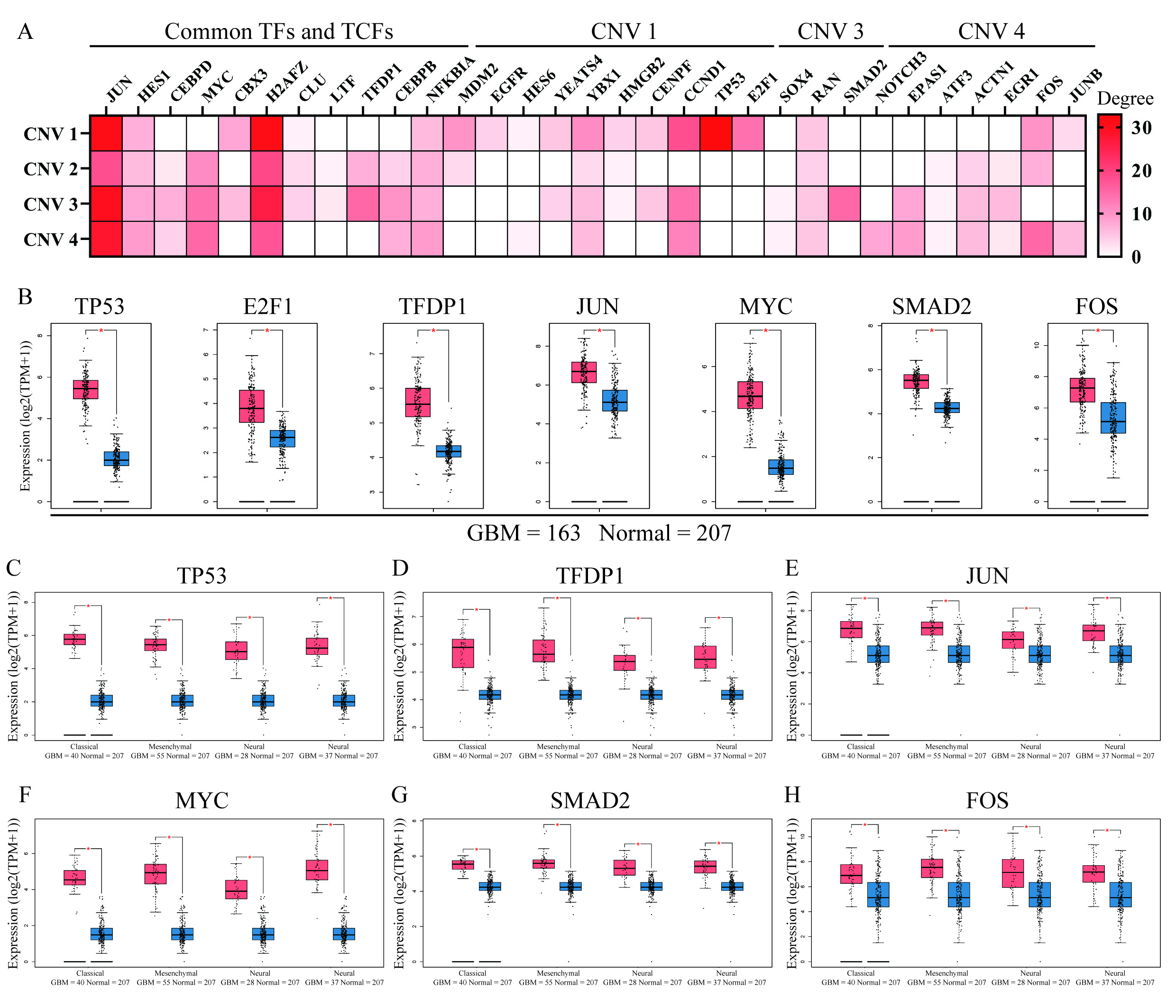

### Supplementary Figure S12

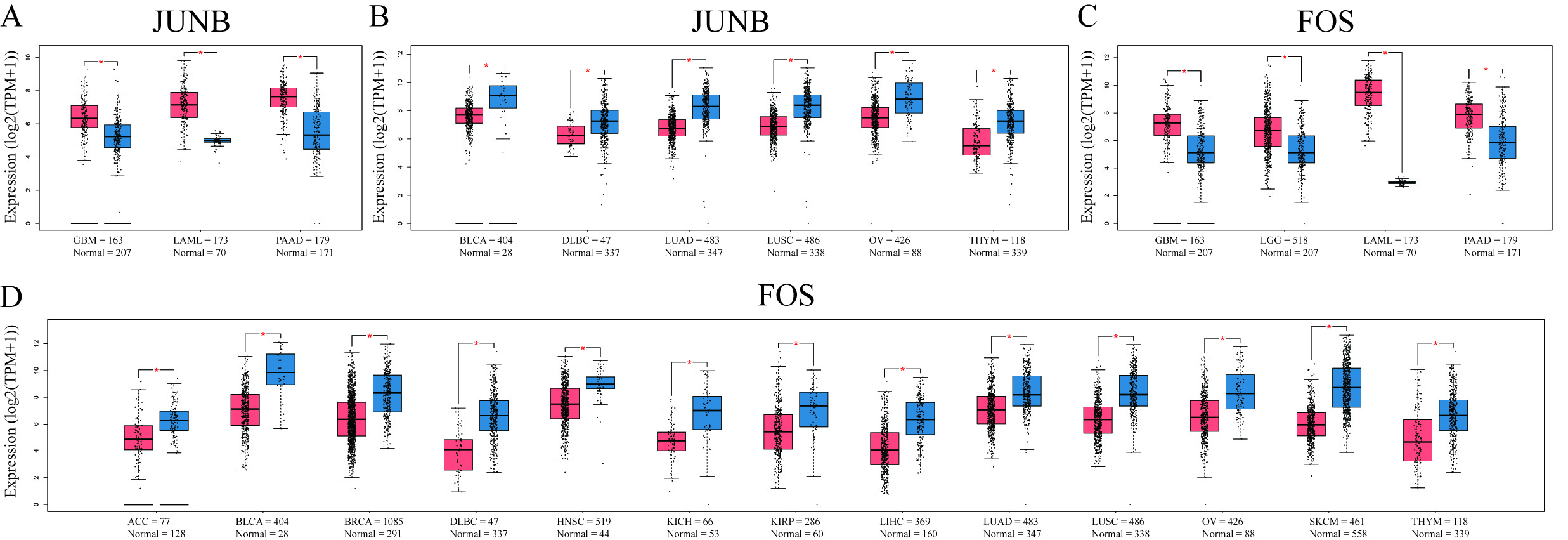

### Supplementary Figure S13

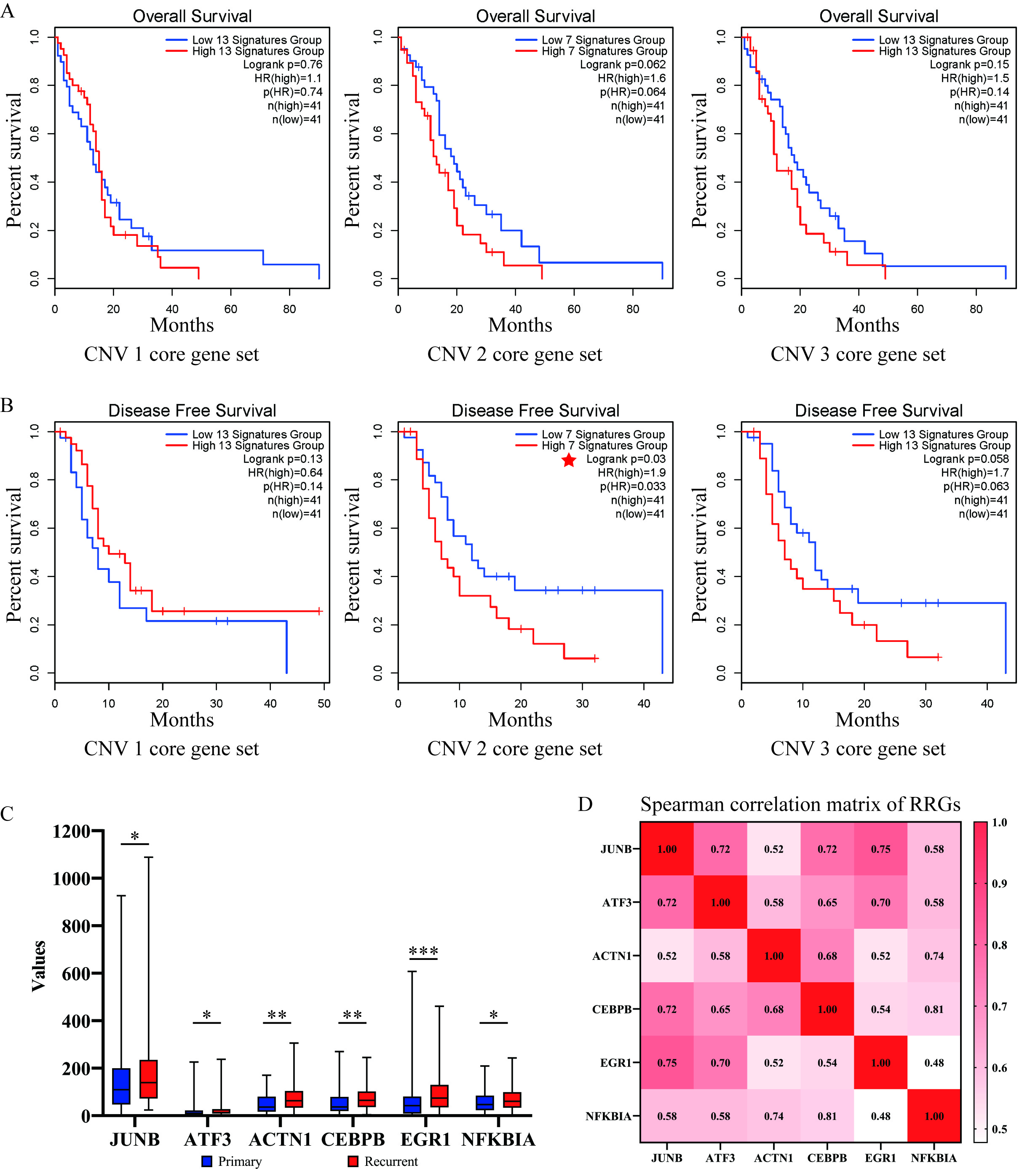

### Supplementary Figure S14

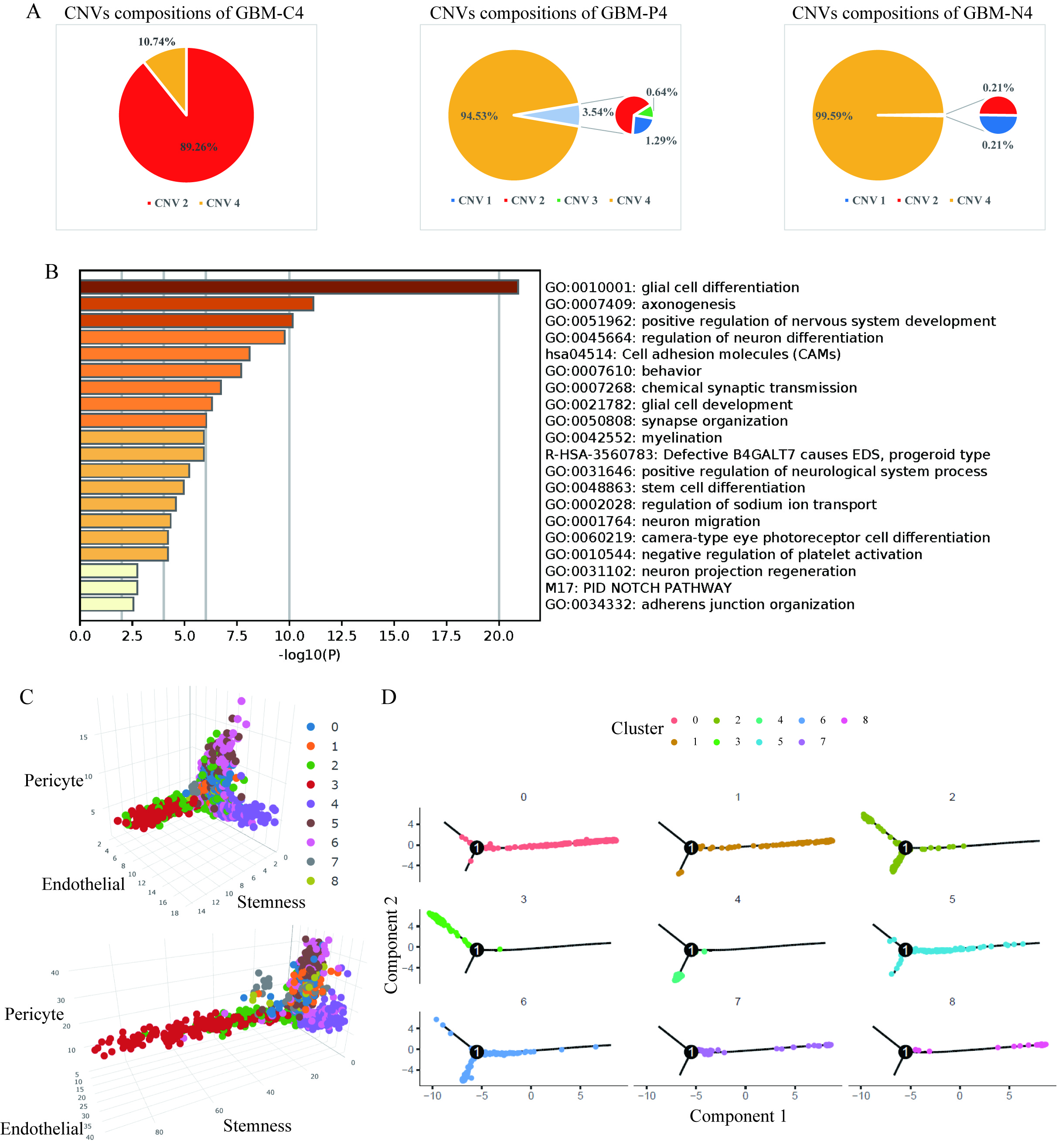

### Supplementary Figure S15

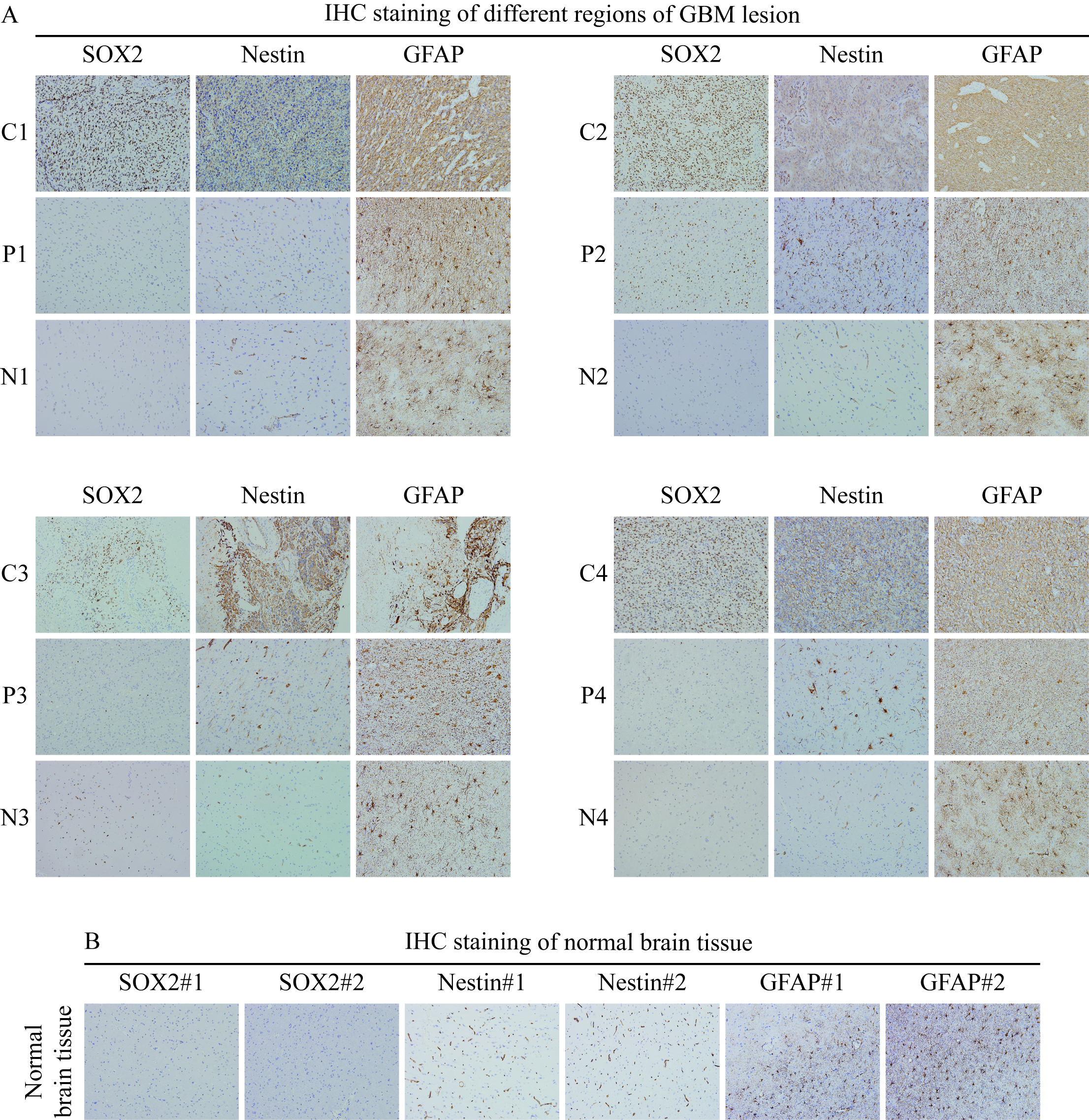

### Supplementary Figure S16

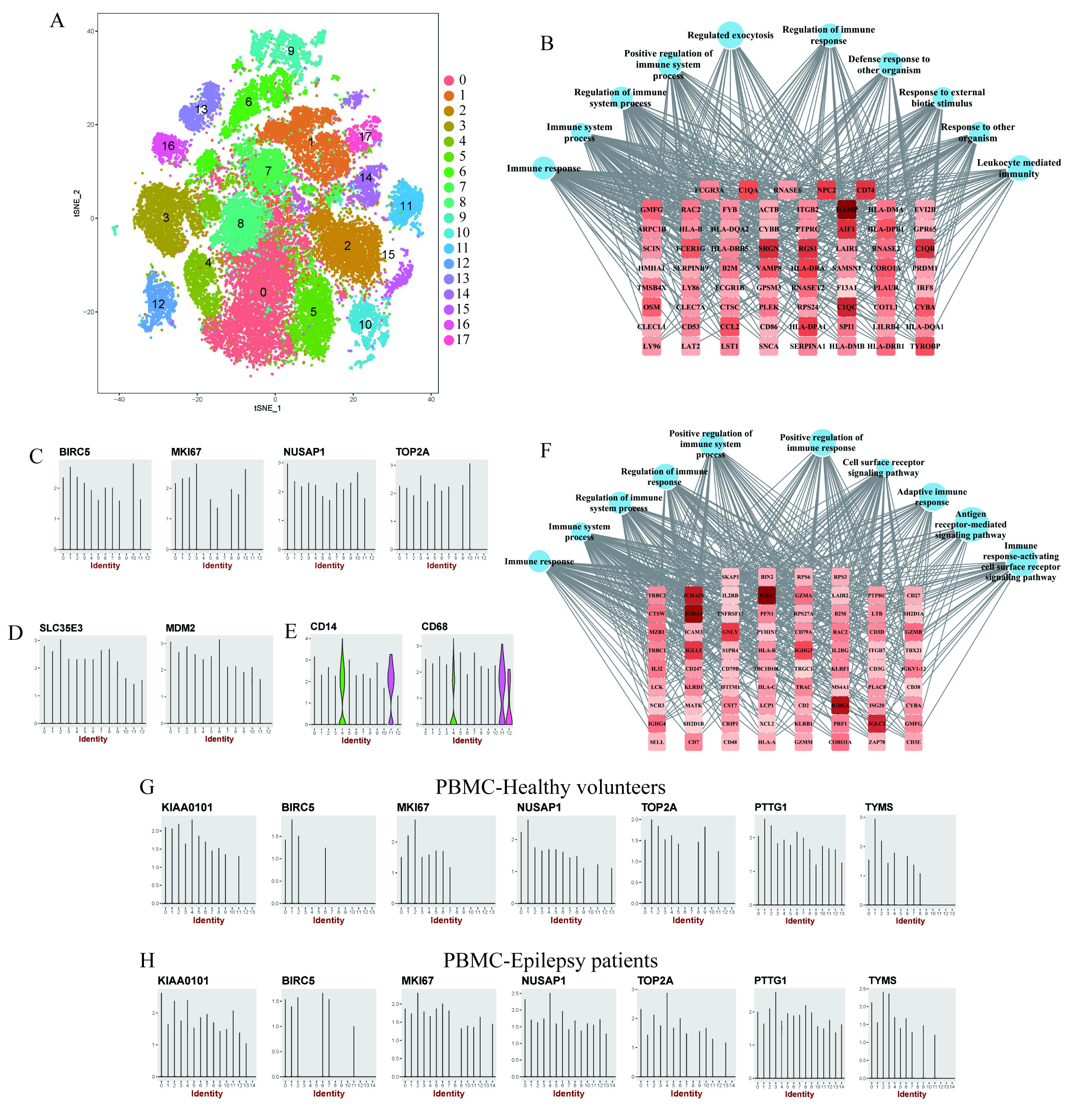
