## Supplementary Table S5 for "Genetic characterization of stem cell-like cancer cell in the peri-tumoral regions and proliferative lymphocyte in the peripheral blood of patients with glioblastoma"

**Supplementary Table S5: Antibodies and DAPI used in this study**

| **Antibodies and DAPI** | **Company** | **Product code** | **Dilution** |
| --- | --- | --- | --- |
| CD133 | Abcam | ab16518 | 1:50 |
| CD34 | Abcam | ab81289 | 1:2000 |
| SOX2 | Abcam | ab79351 | 1:1000 |
| GFAP | Abcam | ab7260 | 1:2000 |
| KI67 | Zhongshanjinqiao | ZM-0167 | Working Concentration |
| Nestin | Zhongshanjinqiao | ZA-0628 | Working Concentration |
| Goat anti-Rabbit (Alexa Fluor 488) preadsorbed | Abcam | ab150081 | 1:500 |
| DAPI | Roche | 10236276001 | 10μM |
